# A Bi-Specific T Cell-Engaging Antibody Triggers Protective Immune Memory and Glioma Microenvironment Remodeling in Immune Competent Preclinical Models

**DOI:** 10.1101/2024.12.18.628714

**Authors:** Markella Zannikou, Joseph T. Duffy, Daniele Procissi, Hinda J. Najem, Rebecca N. Levine, Dolores Hambardzumyan, Catalina Lee-Chang, Lara Leoni, Craig M. Horbinski, Bin Zhang, Amy B Heimberger, Jason M. Miska, Irina V Balyasnikova

## Abstract

**Background:** Bispecific T cell-engagers (BTEs) are engineered antibodies that redirect T cells to target antigen-expressing tumors. BTEs targeting tumor-specific antigens such as interleukin 13 receptor alpha 2 (IL13Rα2) and EGFRvIII have been developed for glioblastoma (GBM). However, there is limited mechanistic understanding of the action of BTE since prior studies were mostly conducted in immunocompromised animal models. To close this gap, the function of BTEs was assessed in the immunosuppressive glioma microenvironment (TME) of orthotopic and genetically engineered mouse models (GEMM) with intact immune systems.

**Methods:** A BTE that bridges CD3 epsilon on murine T cells to IL13Rα2-positive GBM cells was developed and the therapeutic mechanism investigated in immunocompetent mouse models of GBM. Multi-color flow cytometry, single-cell RNA sequencing (scRNA-Seq), multiplex immunofluorescence, and multiparametric magnetic resonance imaging (MRI) across multiple pre-clinical models of GBM were used to evaluate the mechanism and action and response.

**Results:** BTE-mediated interactions between murine T cells and GBM cells triggered T cell activation and antigen-dependent killing of GBM cells. BTE treatment significantly extended the survival of mice bearing IL13Rα2-expressing orthotopic glioma and *de novo* forming GBM in the GEMM. Quantified parametric MR imaging validated the survival data showing a reduction in glioma volume and decreased glioma viability. Flow cytometric and scRNA-seq analyses of the TME revealed robust increases in activated and memory T cells and decreases in immunosuppressive myeloid cells in the brains of mice following BTE treatment.

**Conclusions:** Our data demonstrate that the survival benefits of BTEs in preclinical models of glioma are due to the ability to engage the host immune system in direct killing, induction of immunological memory, and modulation of the TME. These findings provide a deeper insight into the mechanism of BTE actions in GBM.

**WHAT IS ALREADY KNOWN ABOUT THIS TOPIC:** Bi-specific T cell engaging antibodies (BTEs) targeting IL13Rα2 and EGFRvIII have been developed and shown to activate T cells that mediate killing of glioma cells *in vitro* and in i*n vivo*.

**WHAT THIS STUDY ADDS:** By using immune competent preclinical models, this study reveals that BTEs trigger in situ tumor immune memory within the central nervous system.

**HOW THIS STUDY MIGHT AFFECT RESEARCH, PRACTICE OR POLICY:** Insights into the mechanism of action of BTEs inform response biomarkers that should be considered for inclusion in window-of-opportunity clinical trial assessments and for the rational selection of future combinatorial strategies.

## Introduction

Glioblastoma (GBM) is an aggressive and incurable brain tumor with a poor prognosis^1^. The current standard of care for primary disease involves surgical resection, followed by radiation therapy and chemotherapy with the alkylating agent temozolomide^2^. Despite significant advancements in surgery, radiation, and systemic therapy, GBM progression is inevitable. Median overall survival is approximately 15 months, and the 5-year survival rate is less than 5%^3, 4^. At recurrence, there is no established standard of care underscoring the urgent need for novel therapies to enhance survival outcomes in GBM. GBM is regarded as among the most challenging solid tumors to treat due to the blood-tumor barrier limiting the penetration of therapeutics into the glioma microenvironment (TME), immune suppressive TME, heterogeneous expression of tumor-associated antigens, and the limited T cell presence within the tumor^5^.

T cell-based immunotherapeutic treatments in hematopoietic malignancies have been increasingly successful. Among those treatments are bi-specific T cell engagers (BTEs), a class of engineered bispecific antibodies with potent anti-cancer properties. BTEs consist of two single-chain variable fragments (scFvs) derived from different monoclonal antibodies fused by a peptide linker. One part of the scFv of the BTE recognizes a T cell stimulatory receptor such as the CD28 or CD3 epsilon (CD3E) subunit of the T cell receptor complex. The other scFv is specific for a tumor antigen. In bridging T and tumor cells, BTEs activate T cells and direct T cell-mediated killing of malignant cells. Unfortunately, there are significant hurdles in the development of BTE therapy for solid tumors such as GBM. Our group has previously shown the efficacy of a BTE targeting the glioma-associated antigen interleukin 13 receptor subunit alpha 2 (IL13Rα2) in preclinical patient-derived xenograft (PDX) models of GBM. Others have demonstrated a significant therapeutic effect of BTE targeting EGFRvIII, a glioma-specific antigen, in PDX and murine glioma models ^6^, and a clinical study evaluating the safety of BTEs is anticipated (NCT04903795). To overcome the issue of glioma antigen heterogeneity, a DNA-encoded tri-specific T cell engager has been developed to IL12Rα2 and EGFRvIII ^7, 8^. However, these preclinical studies were conducted in immunocompromised hosts lacking a fully functioning immune system rendering them unable to recapitulate the highly immunosuppressive TME in human GBM^9^.

As a potential strategy to overcome the blood-tumor barrier^10^, BTEs with varying target specificities continue to be developed^8, 11–14^. Gaining a deeper understanding of the mechanistic actions of BTEs in fully immunocompetent GBM models is crucial for informing future clinical study designs and rationale selection of combinatorial approaches. Therefore, this study was designed to investigate the therapeutic mechanisms of BTEs in orthotopic and genetically engineered mouse models (GEMM) of GBM. Our findings demonstrate that the activation of murine T cells depends on the binding of BTE to its target, IL13Rα2, both in vitro and in vivo. Moreover, BTE treatment increases the proportion of activated T cells within the glioma and leads to a higher frequency of memory T cells in the gliomas and brains of long-term surviving (LTR) mice. These results suggest that BTE engagement with the host immune system enhances the therapeutic efficacy of BTEs against glioma cells.

## Results

### Generation and functional characterization of the CD3xIL13Rα2 BTE

Single-chain variable fragments derived from the monoclonal antibody 2C11-145 binding murine CD3e^15^ and the monoclonal antibody against human IL13Rα2 (clone 47)^16, 17^ were used to generate a BTE protein suitable for studies in an immunocompetent host. The two scFv were genetically linked with a 23 amino acids GlyS linker^18^. A negative control BTE (NC-BTE) was generated by aligning and replacing the unmatched amino acids in the CDR3 domain of VL scFv targeting IL13Rα2 with those from the scFv p588 sequence - a previously established negative control antibody (MOPC1)^18^. Constructs were generated by subcloning the codon-optimized cDNA encoding BTE and NC-BTE in pLVX-IRES-ZsGreen1 lentiviral constructs (figure 1A). To produce BTE proteins, 293T cells were transduced with BTE and NC-BTE constructs and subsequently flow-sorted to enrich for the brightest ZsGreen1 positive 293T cells. A western blot of the purified BTE confirmed a protein with the anticipated molecular weight of approximately 55 kDa (figure 1B). The specificity of BTE to IL13Rα2 was confirmed with a plate ELISA in which the BTE but not the negative control NC-BTE interacted with recombinant IL13Rα2 (figure 1C). The constructed BTE was specific to IL13Rα2 since no to IL13Rα1 was detected (figure 1C).

**Figure 1.**
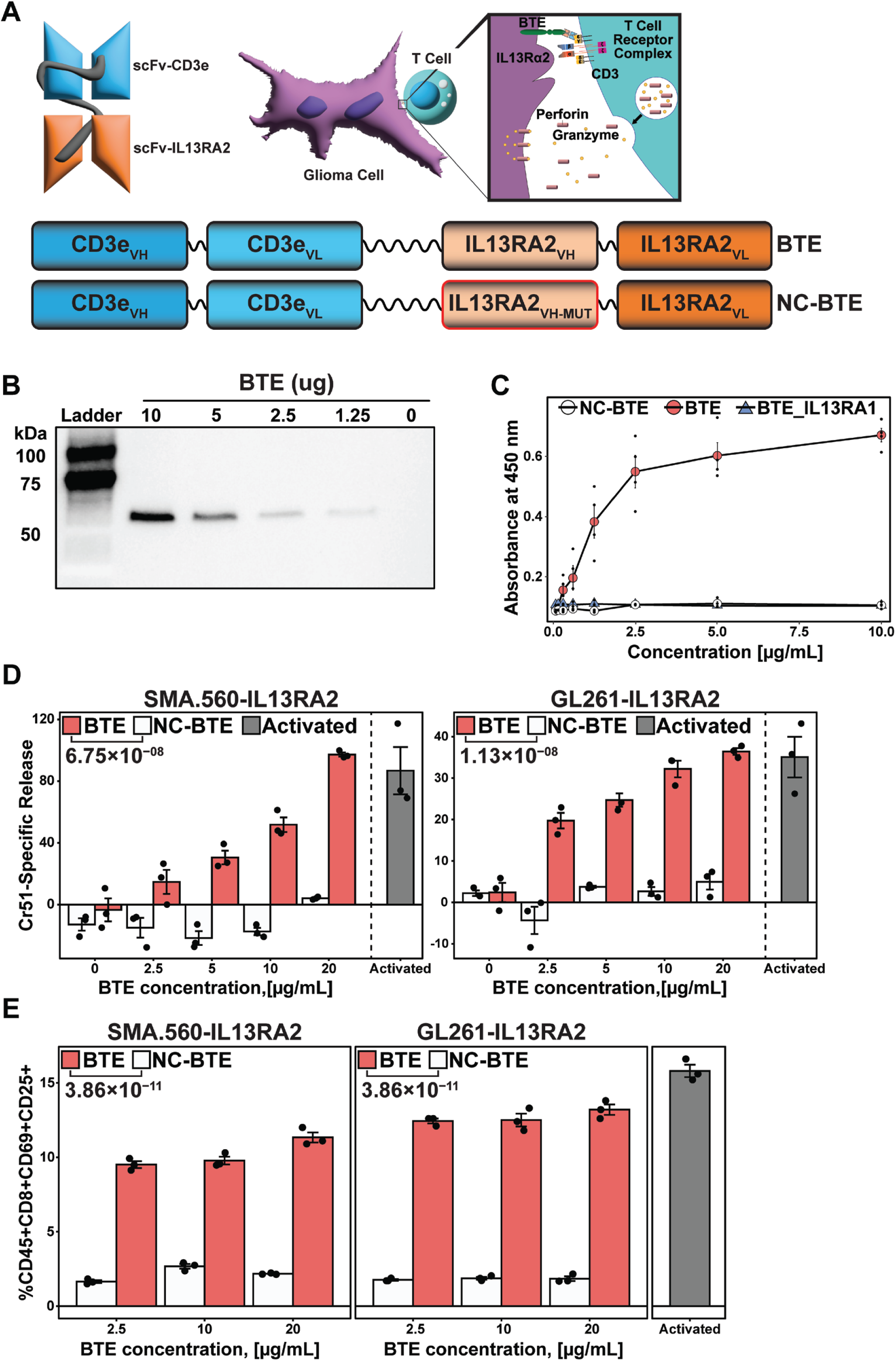
Generation and functional characterization of the CD3xIL13Rα2 BTE. (*A*) Schematic representation of the BTE consisting of the scFv targeting murine CD3E specific component, derived from 2C11-145, and the scFv targeting IL13Rα2, derived from clone47, fused via a flexible GlyS (Gly4Ser) linker. Negative control BTE (NC-BTE) differed from the BTE in the VH domain of IL13Rα2-scFv, which was swapped with the VH domain of scFv p588 of the non-specific MOPC21 antibody. (*B*) A western blot of the affinity-purified BTE at various concentrations was detected using an anti-His HRP conjugated antibody. A single band ∼was observed at 55 kDa. (*C*) ELISA assay showing dose-dependent and specific binding of BTE to recombinant IL13RA2 (red circles), but not to IL13RA1 (blue triangles); NC-BTE (black circles) does not bind to IL13RA2. (*D*) Chromium-51 (Cr51) release assay of murine T cells and glioma cells treated with BTE (red), NC-BTE (white), or Dynabeads™ Mouse T-Activator CD3/CD28 (grey) showing BTE, but not NC-BTE, induced dose-dependent killing of IL13Rα2-expressing SMA560 and GL261 murine glioma cells (Effector-to-target [E:T] ratio=20:1; Incubation time=24 hr; n=3/group). (*E*) Flow cytometric analysis of murine T cells co-cultured with IL13Rα2-expressing SMA560 and GL261 murine glioma cells showing BTE, but not NC-BTE, induced robust expression of T cell activation markers CD69 and CD25 (Effector-to-target [E:T] ratio=20:1; Incubation time = 24 hr; n=3/group). One-way ANOVA followed by Tukey’s Honest Significant Difference (HSD) test for pairwise comparisons)

Next, the ability of BTE to activate T cells and direct T cell-mediated killing of murine glioma cells was evaluated in vitro. The murine glioma cell lines GL261, SMA560, and CT2A previously modified to express IL13Rα2^19^ were used as the targets assessed via Cr^51^-release. T cells activated with CD3/CD28/CD2 immuno-beads served as a positive control. The BTE, but not NC-BTE, significantly increased the killing of SMA560-IL13Rα2 and GL261-IL13Rα2 cells in a dose-dependent manner (figure 1D) but not in the parental SMA.560 and GL261 cells (online supplementary figure 1A). The BTE also triggered significant expression of the early activation makers CD25 and CD69 on CD8+ T cells upon engagement with SMA560-IL13Rα2 and GL261-IL13Rα2 cell lines (figure 1E). This activation was further confirmed with live cell imaging in which increased fluorescent intensity was observed in NOD.Nr4a1GFP/Cre mouse T cells following BTE treatment (online supplementary figure 1B, C). Collectively, these data confirm the creation of a functional BTE and corresponding negative control NC-BTE protein suitable for studies in immunocompetent syngeneic models of glioma.

### BTE treatment in IL13R**α**2-expressing preclinical models enhances survival and expands T cell effector responses in the TME

To clarify if the BTE could be distributed into the brains of GL261-IL13Rα2 glioma-bearing mice, the AF647-labeled BTE was administered intravenously into the tail vein and dynamic live fluorescent imaging was performed longitudinally. The BTE was detected in the brain three hours after administration, with peak signal between 3-6 hours, and diminishing between 24-72 hours (figure 2A). Next, a dose titration at 50 ug/mouse or 200 μg/mouse of the BTE was conducted in SMA.560-IL13Rα2 glioma-bearing mice. These mice were treated with the BTE or saline on 7, 9, and 11 days post-glioma implantation and followed for survival (online supplementary figure 2A). Both concentrations of BTE 50 µg/animal (50 µg/animal: median survival (MS)=18, n=8, p=0.0298 relative to the control; 200 µg/animal: MS=27.5, n=8, p=0.0054 relative to the control) extended the survival of mice compared to saline (MS=15, n=8)(online supplementary figure 2A). No statistical difference in survival was noted between the 50 and 200 µg dose. To confirm the antigen specificity of the BTE, a separate survival experiment was conducted in the parental SMA.560 glioma-bearing mice and there was no increase in survival between the BTE, NC-BTE, and saline group (online supplementary figure 2B). Based on these results, a survival analysis was conducted in the SMA.560-IL13Rα2 glioma-bearing mice treated at 200 ug/animal with either the BTE, NC-BTE, or saline on days 7, 9, and 11 post-glioma implantation (figure 2B). There was no difference in the MS between saline (MS=17, n=7) or the NC-BTE treated group (MS=19, n=8) (figure 2B). The BTE significantly extended the survival of mice (MS=33, n=7, p=0.0035) with 40% surviving over 150 days (figure 2B). Similar survival benefit of the BTE was seen in the CT2A-IL13Rα2 murine glioma model (online supplementary figure 2B).

**Figure 2.**
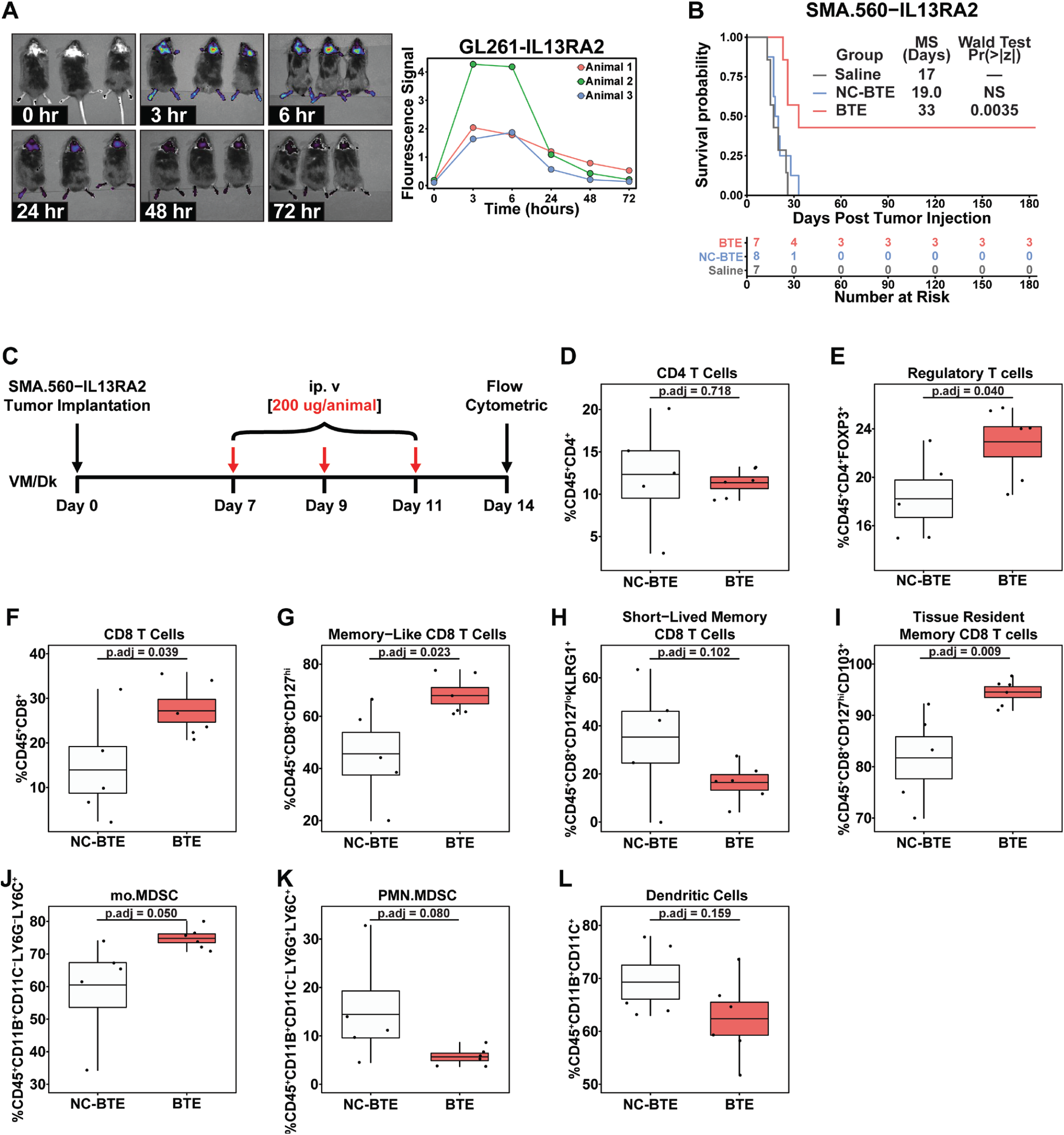
BTE enhances survival and modulates the tumor microenvironment in mice bearing IL13Rα2-expressing gliomas. (*A*) Dynamic live fluorescent imaging of AF647-labeled BTE showing penetration and persistence in the brains of GL261-IL13Rα2 tumor-bearing mice (Fluorescent signal measured as radiance [p/s/cm2/sr]; Image acquisition time = 3, 6, 24, 48, and 72 hrs post AF647-BTE injection; n=3). (*B*) Kaplan–Meier survival curve of SMA.560-IL13Rα2 glioma-bearing mice treated with either BTE (200 µg/animal) (red), NC-BTE (200 µg/animal) (blue), or saline (black). Wald test from Cox proportional hazards regression analysis. (*C*) The experimental schema used for the flow cytometric analysis of the brains of SMA.560-IL13Rα2 glioma-bearing mice following treatment with either BTE or NC-BTE (n=5/group). Flow cytometric analysis revealed (*D*) no change in CD4 helper t cell (CD45+ CD4+), (*E*) increase in regulatory T cell (CD45+ CD4+ FOXP3+), (*F*) increase in CD8 T cell (CD45+ CD8+), (*G*) increase in memory-like CD8 T cell (CD45+CD8+CD127HiKLRG1−), (*H*) no change in short-lived memory CD8 T cells (CD45+ CD8+ CD127Hi KLRG1+), (*I*) increase in tissue-resident memory CD8 T cells (CD45+ CD8+ CD127Hi KLRG1− CD103+), (*J*) increase in monocytic MDSC (mo.MDSC) (CD45+ CD11B+ CD11C−LY6G− LY6C+), (*K*) no change in polymorphonuclear MDSC (PMN.MDSC) (CD45+ CD11B+ CD11C− LY6G+ LY6C+), (*L*) no change in dendritic cells (CD45+ CD11B+ CD11C+). One-way ANOVA followed by Tukey’s Honest Significant Difference (HSD) test for pairwise comparisons).

To determine how the BTE affects the immune composition within the TME, SMA.560-IL13Rα2 glioma-bearing mice were euthanized 3 days after the last dose of either BTE (n=5) or NC-BTE (n=5) (figure 2C). The brains were processed into single-cell suspensions, the cells stained with fluorochrome-conjugated antibodies against immune cell markers, and multi-color flow cytometric analysis was performed. Although there was no difference in the frequency of CD4 T cells between treatment groups (figure 2D), there was an increased frequency of Tregs (CD45^+^CD4^+^FOXP3^+^; p.adj=0.040) (figure 2E), CD8^+^ T cells (CD45^+^CD8^+^; p.adj=0.039) (figure 2F), memory-like CD8^+^ T cells (CD45^+^CD8^+^CD127^hi^; p.adj=0.023) (figure 2G), tissue-resident memory CD8^+^ T cells (CD45^+^CD8^+^CD127^hi^CD103^+^; p.adj=0.009) (figure 2I), and monocytic myeloid-derived suppressor cells (Mo.MDSC)(CD45^+^CD11B^+^CD11C^-^LY6G^-^LY6C^+^; p.adj=0.050) (figure 2J) in the mice treated with BTE relative to NC-BTE. There was no statistical difference in short-lived memory CD8^+^ T cells (CD45^+^CD8^+^CD127^lo^KLRG1^+^; p.adj=0.102) (figure 2H), polymorphonuclear MDSC (Pmn.MDSC)(CD45^+^CD11B^+^CD11C^-^ LY6G^+^LY6C^-^; p.adj=0.080) (figure 2K), or dendritic cells (DC)(CD45^+^CD11B^+^CD11C^+^; p.adj=0.159) (figure 2L).

To further characterize this T cell expansion, both the glioma and spleen from CT2A-IL13Rα2 bearing mice were analyzed with flow cytometry (online supplementary figure 2D). Similar to the findings in the SMA.560-IL13Rα2 model, there was a significant increase in the frequency of Tregs in the brain following BTE treatment (p.adj=0.039), but not in the spleen (online supplementary figure 2E). Markers of T cell cytotoxicity and activation were increased including GZMB (p.adj=0.006) (online supplementary figure 2F), CD69 (p.adj=0.015) (online supplementary figure 2G), and CD25 (p.adj=0.001) (online supplementary figure 2H) in the brain CD8 T cells, but not in the spleen. The brain CD8 T cells upregulated Lag3 expression (p.adj=0.036) (online supplementary figure 2I), but not Tim3 (online supplementary figure 2J) or Pdcd1 (online supplementary figure 2K). Notably, there was an increased frequency of CD44^+^CD62L^-^CD8^+^ effector resident T cells in the brain (p.adj=0.001), but not in the spleen of BTE-treated animals (online supplementary figure 2L, M).

### BTE treatment is protective of glioma rechallenge and generates glioma resident immunological memory

To clarify if the BTE triggers protective immunological memory, SMA.560-IL13Rα2 mice from the prior experiments were rechallenged in the contralateral hemisphere with parental SMA.560 glioma cells (n=5) alongside age-matched control mice (n=5). While age-matched control mice died within 20 days, the SMA.560 rechallenged SMA.560-IL13Rα2 mice survived over 210 days without neurological symptoms (figure 3A). These long-term survivors (LTS) were then euthanized and the brains were examined by histology (n=2/group) and flow cytometry (n=3/group). H&E staining revealed the presence of gliomas in control mice that died within 20 days but the absence of gliomas in the LTS (figure 3B). Immunohistochemical (IHC) staining of CD8 on serial tissue sections revealed a wide distribution of these cells throughout the brain of rechallenged LTS mice (figure 3C). Quantitative analysis showed significantly more CD8+ T cells in the brains of LTS compared to controls in both the ipsilateral (p.adj=2.22×10−16) and contralateral (p.adj=4.78×10−7) hemispheres of the glioma implantation site (figure 3D). In the flow cytometric analysis of the LTS and control brains, there was a significant increase in the frequency of CD8 effector resident T cells (CD44^+^CD62L^-^) (p.adj=0.0035) (figure 3F) and tissue-resident memory CD8 T cells (CD44^+^ CD62L^-^CD103^+^) (p.adj=0.028) (figure 3G) in the brains of rechallenged LTS mice compared to control. Moreover, Ifng and TNFα expression was significantly increased in the CD8+ T cells of the LTS compared to control (p.adj=0.093) (figure 3G). Gliomas are known to be highly infiltrated by CD11b^+^ cells. There was a significantly lower frequency of CD11b^+^ cells in the brain of LTS mice compared to the control (figure 3I) (p.adj=0.001).

**Figure 3.**
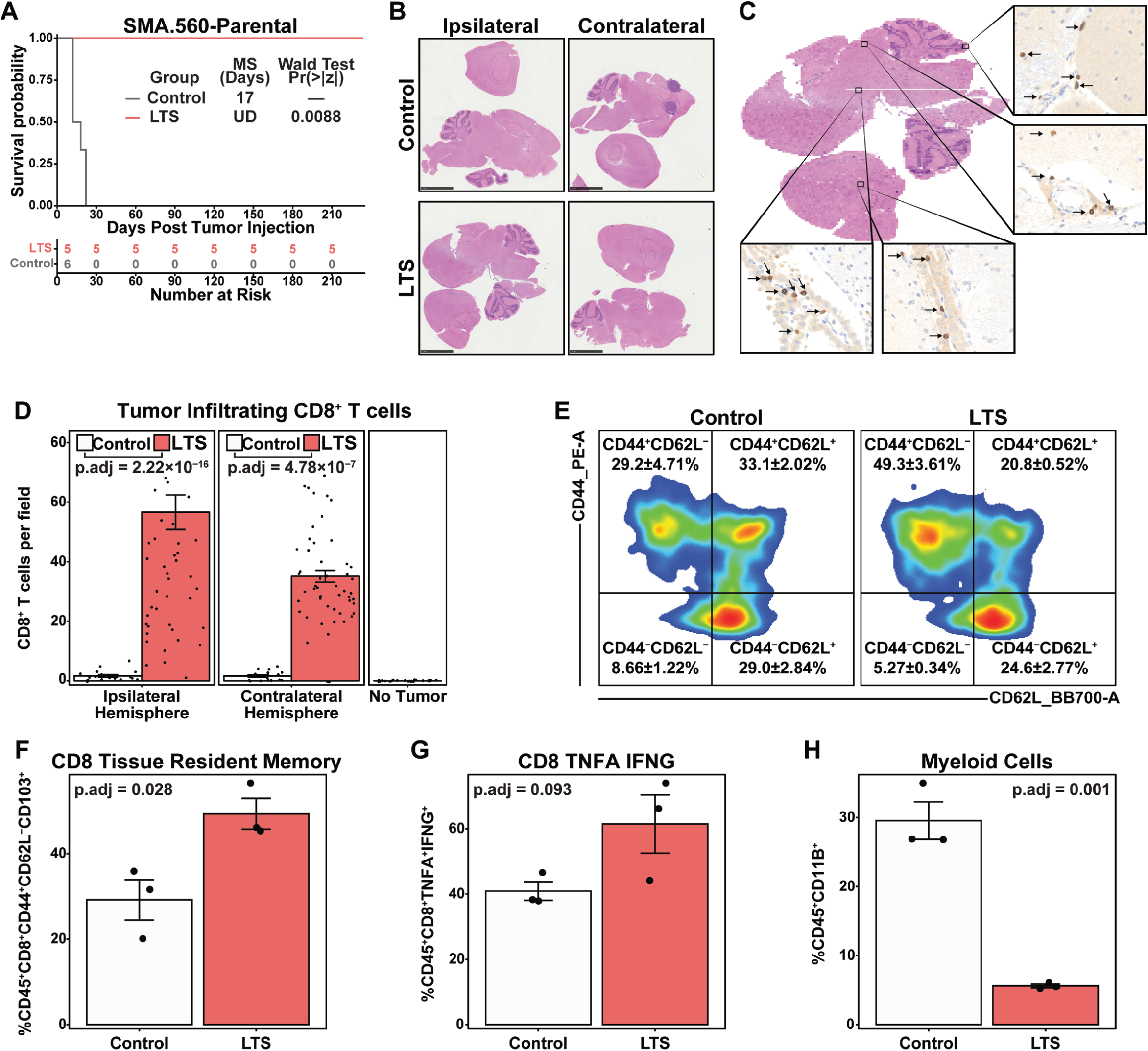
The rechallenge of LTS mice with gliomas demonstrated the generation of memory. (*A*) Kaplan–Meier survival curve of long-term surviving (LTS) (red) and age-matched control (black) mice following implantation of parental SMA.560 murine gliomas into the contralateral brain hemisphere to the primary tumor implantation site. Animals were followed for survival, and statistical significance was tested via the Wald test from Cox proportional hazards regression analysis. (*B*) The mice of LTS (210 days post-secondary injection) and age-matched control (at endpoint) were sacrificed and stained using H&E to evaluate for the presence of tumors in the ipsilateral and contralateral hemispheres of the implantation site. Scale bar represents 2.5 mm. (*C, D*) Serial sections of the same brains stained for infiltrating CD8+ T cells. (*C*) Representative image of one LTS-ipsilateral section with magnified images in the cerebrum (arrowheads) and the cerebellum (arrowheads). The scale bar represents 2.5 mm for the H&E image and 100 µm for magnified anti-CD8 IHC. (*D*) Quantitative analysis of CD8+ T cells in the ipsilateral and contralateral hemispheres of the brain of LTS, age-matched control, and non-tumor bearing mice. One-way ANOVA. (*E-H*) Flow cytometric analysis of freshly dissected brains of LTS (red) and age-matched control (white); n=3/group. (*E*) A representative contour plot shows an increased frequency of CD8 resident T cells (CD44+CD62L-) in the brains of LTS (left) compared to the control (right). Quantified frequency of (F) tissue-resident memory CD8 T cells (CD8+ CD44+ CD62L-CD103+), (G) inflammatory cytokine producing CD8 T cell (CD8+ TNFα +IFNγ +), and (H) total population of myeloid cells (CD45+ CD11B+). Differences among multiple groups were evaluated using one-way or two-way ANOVA with Tukey’s Honest Significant Difference (HSD) test.

### BTEs have a therapeutic effect in high-grade GEMMs

To more closely model human GBM, the BTE was evaluated in a GEMM with de novo gliomagenesis in adult mice ^20^. The GEMM generates IL13Rα2-expressing gliomas with loss of p53 and PTEN and overexpression of PDGFB. Histological analysis of GEMM gliomas revealed key morphological characteristics of human GBM including pseudo palisading necrosis, microvascular proliferation, hemorrhage, heterogeneous expression of IL13Rα2, infiltration of CD11b^+^ macrophages, and a low frequency of CD3^+^ T cells within tumor bed (figure 4A). Treatment of the GEMM with the BTE (n=20, MS=50.5 days) significantly extended survival compared to control (n=19, MS=41 days) (p.adj=0.0045) (figure 4B). To further characterize the changes within TME induced by the BTE, cells from the glioma and spleen were analyzed via flow cytometry 48 hours after the final treatment (figure 4C). In this model, there was no difference in Tregs in the brain between the BTE and control group (CD45^+^CD4^+^FOXP3^+^) (p.adj=0.613) (Fig. 4D). Similar to the findings in the implanted orthotopic gliomas, there was increased expression of the CD69 activation marker in the brain but not the spleen of the BTE-treated animals (p.adj=0.016) (figure 4E). In contrast, there was no difference in the GZMB cytotoxicity marker GZMB (p.adj=0.542) (figure 4F), possibly related to the kinetic differences of expression secondary to the time point of sampling between the models. There was an increase in tissue-resident memory CD8 T cells (CD44^+^ CD62L^-^CD103^+^) (p.adj=0.022) (figure 4G) and effector resident CD8 T cells (CD44^+^ CD62L^-^) (figure 4H). Lastly, multiplex seqIF visualization of the TME of the GEMM treated with BTE and saline revealed that the IL13Rα2 (green) expression is retained in tumor tissue after 3 weeks of BTE treatment (figure 4I).

**Figure 4.**
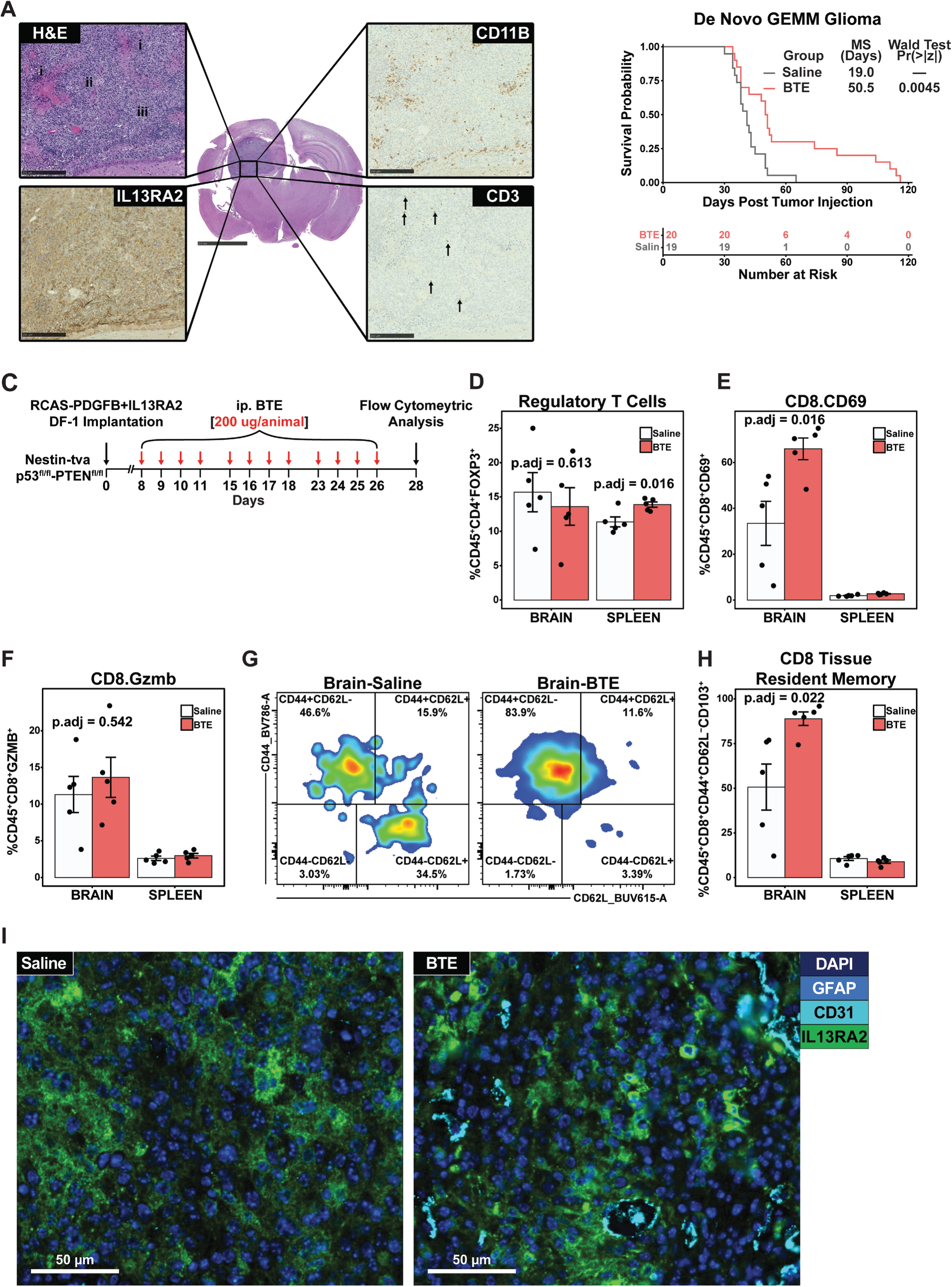
BTE treatment results in therapeutic benefits in genetically engineered mice bearing de novo glioma tumors. (*A*) Histopathological analysis of RCAS-P53fl/fl-PTENfl/fl-IL13Rα2 GEMM glioma model with high magnification images showing morphological characteristics similar to human GBM. H&E (top-left) shows areas of pseudopalisading necrosis (i), microvascular proliferation (ii), and hyperplasia (iii). IHC staining revealed many infiltrating CD11B+ macrophages (top-right), scarce infiltrating CD3+ T cells (bottom-right), and heterogeneous IL13Rα2 expression (bottom-left) within the tumor bed. (*B*) Kaplan–Meier survival curve of the de novo GEMM treated with BTE (red) (n=20) or saline (black) (n=19). Animals were followed for survival, and statistical significance was tested via Wald test from Cox proportional hazards regression analysis. (*C*) An experimental setup was used in flow cytometric analysis of the brains and spleens of the GEMM following treatment with either BTE or saline (n=5/group). Flow cytometric analysis revealed (*D*) no change in regulatory T cell (CD45+ CD4+ FOXP3+) and brain-specific increases in (*E*) activated CD8 T cell (CD45+ CD8+ CD69+), (*F*) cytotoxic CD8 T cell (CD45+ CD8+ CD69+), (*H*) resident CD8 T cells (CD45+ CD8+ CD44+ CD62L-), and (*G*) tissue-resident memory CD8 T cells (CD45+ CD8+ CD44+ CD62L-CD103+). (*I*) Multiplex IF of saline (left) and the BTE (right) treated GEMM demonstrating that there was no difference in IL13Rα2 expression. Visualization of tumor architecture was done via staining of the nuclei with DAPI, glioma cells with GFAP, and blood vessels with CD31. Scale bars equal to 50µm

### BTE treatment remodels the TME to immune effector responses in de novo gliomas

To gain insights into how the BTE impacts the TME, scRNA-seq was performed in the GEMM model following BTE treatment. Like human GBM, the GEMM TME is dominated by the resident and peripheral myeloid cells (figure 5A). UMAP analysis as a function of treatment showed less peripheral myeloid cells in the BTE-treated animals compared to saline treatment (figure 5B) likely due to the reduced glioma size in the BTE-treated animals. After sub-clustering and performing dimensional reduction on the T and NK cell populations (online supplementary figure 3A), these cells showed distinct differences in gene ontology based on treatment (figure 5C). Notably, the relative frequency of these cells remained similar between treatment groups (online supplementary figure 3B). Subsequent gene ontology (GO) enrichment analysis of significant differential gene expression (DEG) mapped to pathways involved in cytokine production, T cell differentiation, T cell activation, and antigen processing and presentation in the CD8 T cell cluster in the BTE-treated group (figure 5C). Evaluation of specific genes key to CD8 T cell effector function revealed elevated expression of genes involved in activation, cytotoxicity, and memory with reduced expression of exhaustion genes (figure 5D). Memory and effector function findings, such as CD8 and CD4 T cells co-expressing TCF1 (figure 5E) and IFNγ (figure 5F), were validated with multiplex immunofluorescence visualizing the TME in the GEMM treated with BTE or saline.

**Figure 5.**
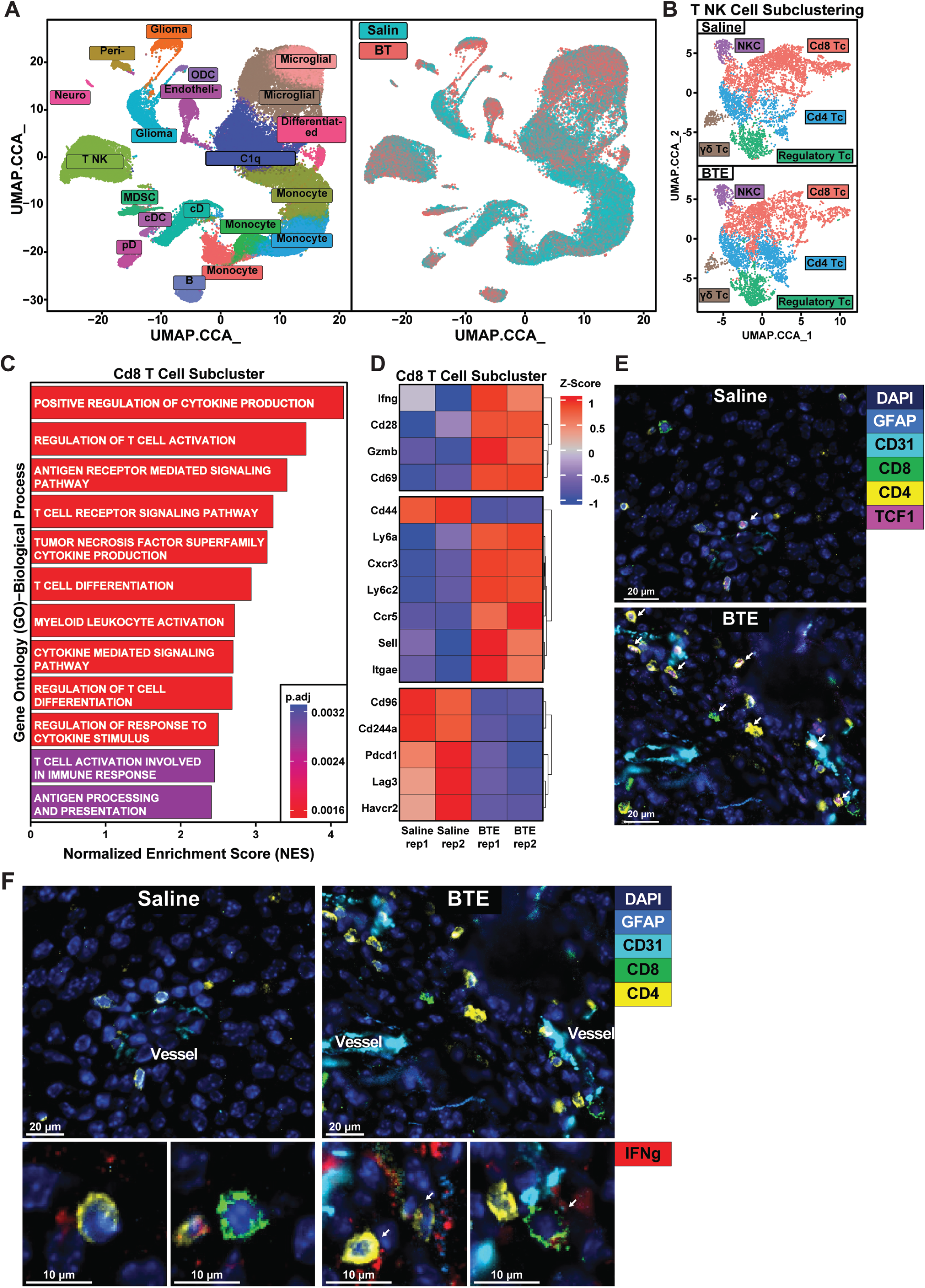
Sc-RNA seq reveals remodeling of tumor-microenvironment in de novo glioma tumors upon BTE treatment. (*A*) UMAP visualization of single-cell RNA sequencing data from cells isolated from the tumors of de novo GEM glioma models. The data were derived from two independent experiments, each with two treatment groups (n=5/group): BTE-treated and saline-treated. In each experiment, isolated cells from each mouse were pooled together for each group, resulting in a total of four samples (two BTE-treated and two saline-treated) totaling to 64,124 cells. Canonical Correlation Analysis (CCA) integration was used to combine data from the two experiments into a shared UMAP, with the left panel colored by cell cluster identity and the right panel colored by treatment group. Cells were enriched for CD45-positive populations at a CD45+:CD45-ratio of 8:2. (*B*) UMAP visualization of T and NK cell subclusters following CCA integration displayed based on saline (top) or BTE treatment (bottom). Subclusters include CD8 T cells (red), CD4 T cells (blue), regulatory T cells (green), γδ T cells (brown), and NK cells (purple). (*C*) Gene ontology analysis of differentially expressed genes (DEGs) in CD8 T cells that were upregulated in BTE treated samples compared to Saline. (*D*) Heatmap of DEGs in CD8 T cell cluster across all samples related to activation-cytotoxicity, CD8 T cell memory, and exhaustion as a function of treatment. (*E*) Multiplex IF imaging of the gliomas in the GEMM model treated with saline (top) or the BTE (bottom). Visualization of tumor architecture was done via staining of the nuclei with DAPI, glioma cells with GFAP, and blood vessels with CD31. High magnification images show glioma infiltration with CD8 (green) and CD4 (yellow) T cells with intracellular TCF1 (magenta) following BTE treatment denoted with single arrows; Scale bars: 20µm. (*F)* Multiplex immunofluorescence imaging of the gliomas in the GEMM model treated with saline (top-left) or the BTE (top-right) Sections were stained for tumor architecture DAPI, GFAP, and CD31, and T cell subsets with CD8 (green) and CD4 (yellow). Below each image are high magnification images of glioma infiltrated CD8 (bottom-left) and CD4 (bottom-right) T cells with intracellular IFNγ (red). Scale bars for bigger panels: 20µm, for smaller panels: 10µm.

### Volumetric MRI reveals tumor-specific radiographic responses and increased TME complexity after BTE treatment

To gain deeper insights into the therapeutic effects of BTEs on tumor progression, therapeutic response, and TME changes in the GEMM, advanced MRI techniques were utilized. Representative 2D MRI images of the glioma (circled in yellow) and the corresponding 3D rendered glioma image generated following manual segmentation across the whole brain is shown in figure 6A. BTE treatment relative to saline reduced the glioma tumor volume (online supplementary figure 3C). In addition to measuring tumor size, T2 parametric tumor maps were generated from the MRI 2D images to quantify tumor viability and therapy response. The intra-tumor T2 values of MR 2D images from BTE- and saline-treated animals were visualized by superimposing quantitative, color-coded parametric tumor maps. These maps were generated from multi-echo sequences using voxel-by-voxel least-squares fitting procedures (figure 6B). Further color-coded (red to blue) parametric mapping of 2D MR images in multi-voxel regions revealed specific differences in radiomic-like features and patterns between BTE and saline-treated animals (figure 6C). Derivation of quantitative tumor response and tumor viability indexes from the combination of these in vivo indexes revealed significantly higher tumor response indexes in BTE-treated animals compared to saline controls (figure 6D).

**Figure. 6:**
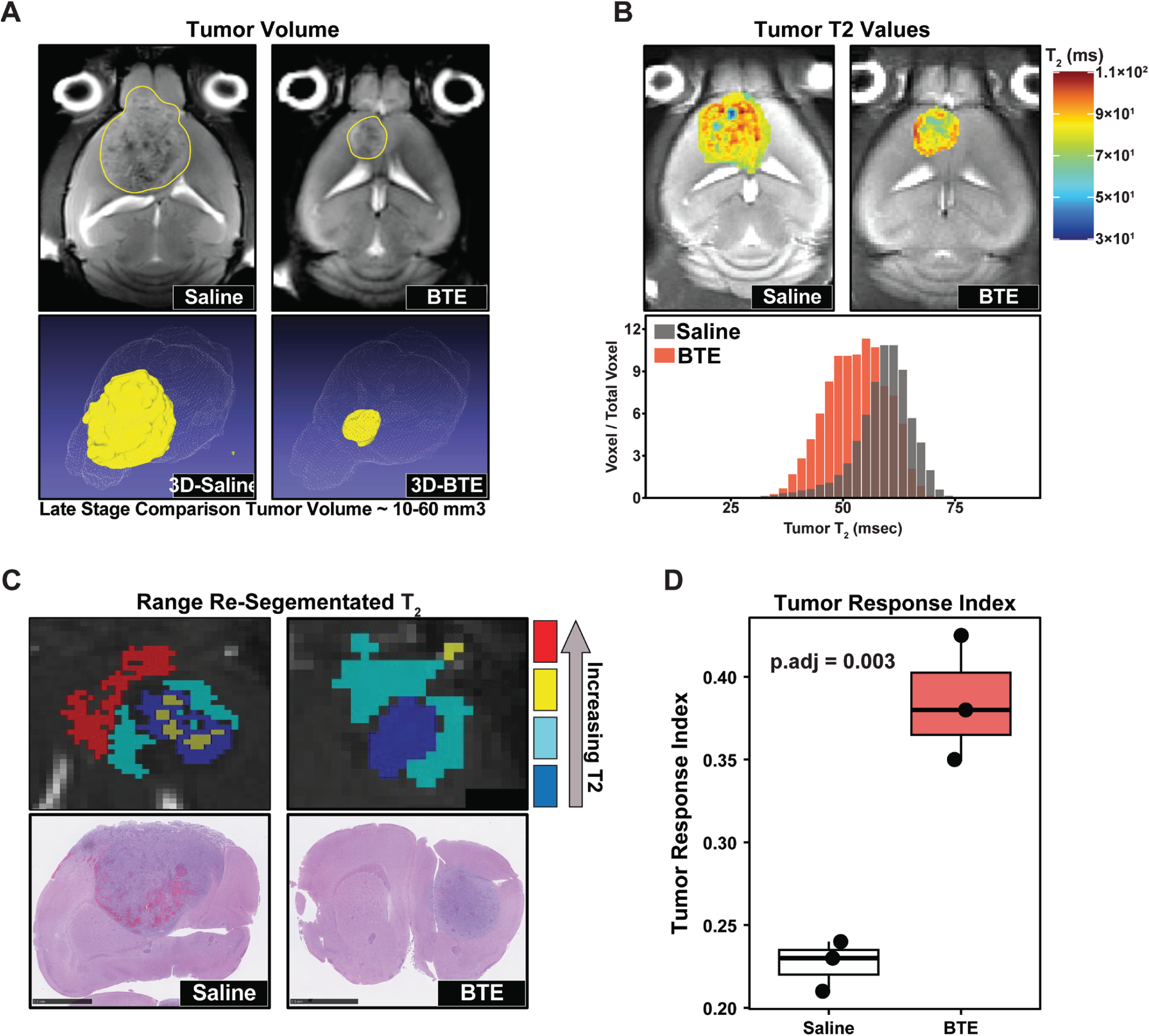
Magnetic Resonance Imaging demonstrates responses of de novo tumors to BTE in GEMM. (*A*) Representative image set of two MRI gray-scale anatomical brain sections from saline control (left) and BTE-treated (right) mice. Whole-tumor 3D delineation was done to extract tumor size and to enable rendered 3D visualization of the overall tumor size. (*B*) A representative set of two quantitative T2 parametric colored maps for a saline control and BTE-treated mouse overlayed on a corresponding anatomical gray-scale image. (*C*) Comparative clustered T2 parametric image of each tumor (control saline on the left and BTE-treated on the right) with different colors depicting intratumor regions with different T2 values (BLUE/CYAN low values and YELLOW/RED high values). An H&E stain of a representative section for each mouse was selected to match the exact location of the MRI-derived image as closely as possible (using anatomical features). The clustered intra-tumor regions were then used to extract volumetric quantities for each T2 range, with the driving hypothesis (from our work and existing literature) that an overall decrease in T2 tumor values can be associated with therapeutic response. (*D*) Box-chart plot of each Tumor Response Index value in each mouse, which agrees with survival analysis, flow cytometry, and RNAseq data.

## Discussion

BTEs are recognized as potent targeted T cell-based immune therapeutics that function independently of MHC-restricted antigen recognition and exhibit strong anti-tumor activity^21, 22^. This type of therapeutic strategy is highly relevant for solid cancers such as GBM, which downregulates MHC expression as an immune evasion strategy. Since the FDA approval of Blinatumomab in 2014 for B-cell precursor acute lymphoblastic leukemia^23^, eight BTEs have been approved for use against hematopoietic malignancies. Driven by this clinical success, over 100 distinct BTEs have been evaluated in more than 250 clinical trials for a variety of applications in solid tumors^24^. However, the complex TME in solid tumors poses unique challenges for BTE therapies not encountered in liquid malignancies, and only two BTEs, Tarlatamab (small cell lung cancer)^25^ and Tebentafusp (melanoma)^26^, have been FDA-approved. In GBM, the highly immunosuppressive, myeloid cell-dominated TME thwarts anti-tumor immune activity^27–29^. Our group and others have demonstrated that BTEs targeting IL13Rα2 and EGFRvIII elicit a powerful therapeutic effect in PDX mouse models of GBM ^7, 8, 18, 19^. Yet, the immunodeficient mice used in these studies fail to recapitulate the hallmark immunosuppressive TME of GBM. Thus, investigating how BTEs behave in GBM models with an immune intact TME is critical for understanding their clinically relevant therapeutic potential.

To generate the BTE suitable for studies in an immunocompetent host, we genetically fused a scFv145-2C11 recognizing murine CD3e with our anti-IL13Rα2 scFv via flexible linker based on our previously validated BTE configuration^18^. *In vitro*, the resulting BTE activated and directed murine T cell-mediated killing of IL13Rα2-positive murine glioma cells. *In vivo*, BTE accumulated in the brains of immunocompetent mice bearing IL13Rα2-positive murine glioma after systemic injection. As such, we hypothesized that BTEs would engage host T cells and glioma cells in the brain and retain anti-tumor activity in immunocompetent models of GBM. Consistent with our hypothesis, systemic delivery of BTE resulted in target-specific T cell activation confined to the brain tumor, with no activation observed in peripheral tissues. We demonstrated that the BTE exhibited anti-tumor activity, extending survival across three murine GBM models. The therapeutic effect was target-dependent, as no response was seen in BTE-treated mice bearing parental glioma tumors lacking the IL13Rα2. Unlike treatments with CAR T cells ^30, 31^, no loss of IL13Rα2 expression in tumor tissue was observed after 3 cycles of BTE treatments in GEMM at this time point. This specific on-cancer activation of T cells observed in this study is of importance. Using BTEs when the target is expressed in healthy tissues, even at low levels, can lead to significant damage to normal organs and neurotoxicity due to inadvertent T cell activation distant from the tumor. In fact, BTE-induced on-target, off-tumor toxicities are well-documented in solid tumors and warrant careful consideration^32, 33^. In that respect, it is well established that IL13Rα2 is a glioma-specific antigen ^34–36^ and expressed in other peripheral cancers^37–42^ and brain metastases^43^, but not normal tissues. The specific expression of IL13Rα2 in glioma but not in other cells provides reassurance of the safety of the BTE, which will nonetheless be evaluated in future toxicity studies using a humanized BTE protein.

The scarcity of antigens expressed in GBM makes IL13Rα2 and EGFRvIII, two antigens expressed exclusively in glioma cells and not in normal brain tissue, unique and suitable for targeted applications. Not surprisingly, these two targets are the most tested in pre-clinical and clinical settings for GBM^44^. BTE targeting EGFRvIII has been shown to significantly improve the survival of mice bearing human orthotopic glioma tumors^6^ and a clinical study evaluating the safety of this BTE is anticipated (NCT04903795). Our group has previously demonstrated that BTE-secreting neural stem cells improve the survival of mice bearing patient-derived glioma xenografts expressing IL13Rα2^18^. Others have also reported the therapeutic effect of different BTE configurations in GBM as a gene therapy approach^45^. However, the immense inter- and intratumoral antigen heterogeneity in GBM presents major challenges for targeted therapies, as targeting a single antigen often leads to antigen escape ^30, 46^. Park and co-authors addressed this limitation using a gene therapy approach. They co-delivered two DNA-encoded BTEs targeting EGFRvIII and HER2, demonstrating superior tumor growth suppression and clearance compared to either BTE alone^7, 8^. Others adopted a different approach to overcome GBM heterogeneity by creating multi-specific therapies, such as EGFRvIII-CAR T cells secreting anti-EGFR BTEs ^12^ or employing EGFR-IL13Rα2-specific CAR T cells^11^. Both of these studies showed superior efficacy compared to non-BTE secreting CAR T cells in immunocompromised mice bearing orthotopic GBM. Two clinical trials (<u>NCT05168423</u>, <u>NCT05660369</u>) evaluating the safety and feasibility of CART-EGFR-IL13Rα2 cells^13^ and a EGFRvIII CAR-T cell secreting a EGFR-specific BTE ^14^ have been initiated.

The studies we conducted in immunocompetent hosts not only determined the ability of BTE to activate T cells in the immunosuppressive TME of GBM but also revealed additional mechanisms contributing to the therapeutic action of BTEs. For example, the analysis of tissues revealed a significant enrichment of TRM T cells in the brains of tumor-bearing mice treated with BTE across the GBM models evaluated. The increased frequency of these cells likely contributed to the glioma rejection in the LTS mice rechallenged with the parental glioma line lacking IL13Rα2 expression suggesting induced epitope spreading. Notably, the GEMM, which more closely recapitulates the morphology of human GBM including a low frequency of T cells, also responded to BTE treatment and showed an increased frequency of TRM CD8 T cells. The scRNA sequencing in this GEMM showed that the CD8 T cells in the brains of BTE-treated animals had enhanced cytotoxic (CD69, GZMB, CD28, and IFNγ) and memory (Sell, CD44, Itgae, CxCr3, LY6C2, CCR5) profiles but reduced exhaustion profiles (Lag3, Havcr2, Pdcd1, CC244a, and CD96) compared to control mice. The presence of Tcf1-positive T cells was greatly elevated in tumor tissue from BTE-treated mice compared to the control. Tcf1 has been shown to play an important role in T cells development and memory, including responses to cancer^47^. TRM T cells are known to stay localized in peripheral tissues, continuously monitor the surroundings, and remain primed to rapidly respond to distress signals. These traits position TRM T cells as key contributors to anti-tumor surveillance. There is evidence supporting that the effectiveness of immunotherapy relies on the generation of TRM T cells^48^. In glioma, the presence of TRM T cells was reported in context of T-αFGL2 cell treatment, peptide and virus treatments ^49–51^. Virus-specific memory T cells have been found to populate the mouse and human GBM TME and could be re-activated in the experimental setting. Moreover, a high presence of CD8^+^ TRM cells has been shown to correlate with better survival in glioma^52^. Evaluation of tumor tissue from both lung^53^ and melanoma cancer patients^54^ treated with anti-PD-1 and responded, had increases in TRM CD8 T cells frequency. In patients who were treated with CAR-T cell therapy, enriched populations of TRM T cells were seen in responders to CD19- and BCMA-specific CAR T-cells^55^. Given the established importance of TRM T cells to immunotherapeutic response and sustained anti-cancer immunity, the observation that TRM cells may have formed memory against glioma cells likely participated in the therapeutic response to rechallenge in the BTE-treated group.

Non-invasive monitoring of GBM, especially in the context of immunotherapy to differentiate between tumor progression and treatment-related neuroinflammation, has become a major focus of study in recent years^56–58^. Advanced MRI imaging techniques were used in this study to better understand BTE therapeutic effects on tumor dynamics in the GEMM. Qualitative visualization of non-invasive MRI tumor images exhibited both size and T2 quantitative features and patterns consistent with a therapeutic effect of BTE. In addition to a clear reduction in average tumor size detected through MRI-based volumetrics, the BTE group also exhibited less heterogeneous T2 patterns with overall decreased T2 values relative to saline-treated tumors. These changes may be associated with alterations in the TME cell composition as quantified by flow cytometric analyses and scRNA sequencing analysis of the gliomas. It has been shown that the heterogeneity and higher T2 values correspond to a more complex microenvironment associated with rapidly progressing tumors^59–61^. Despite the limitations of the analysis, the multiparametric MRI analysis could provide a non-invasive response assessment in the future.

Studies in immunocompetent hosts provide needed insights into the behavior of T cells. However, there are well-documented differences between mouse and human T cells^62^. The extracellular domains of CD3E exhibit a low level of amino acid sequence conservation (47%) between humans and mice^63^. The BTE we designed and tested in this study binds murine, but not human T cells^64^. It has been shown that BTEs also exhibit a relatively short plasma half-life^65^. In that respect, several modifications of BTE, such as fusion with engineered Fc or human serum albumin, offer the opportunity to significantly expand the time of the BTE in systemic circulation^66–69^. In the context of brain tumors, the balance between the circulation time and the size of the protein should be carefully investigated to ensure sufficient penetration in GBM tissue. Thus, design of reagents suitable for both human and mouse studies, along with modifications that enhance penetration into brain tumors and improve plasma stability, will be essential to overcome the limitations of pre-clinical testing for the clinical translation of BTEs.

In conclusion, our study demonstrates the survival benefits of BTEs in several immune-competent models of GBM, along with specific immune engagement with the target, induction of immunological memory, and modulation of the TME. This provides strong evidence of BTE-induced responses and the engagement of the host immune system, reinforcing the mechanism of action. These findings also encourage further investigation into other brain and peripheral cancers expressing IL13Rα2. While combating GBM remains our primary focus, translating this immunotherapeutic strategy may offer therapeutic benefits in other solid tumors, warranting further exploration of its full potential.

## Methods

### Cell culture

GL261-IL13Rα2, CT2A-IL13Rα2, and SMA-560-IL13Rv2 were modified with human IL13Rα2 as previously described^19^. Glioma cell lines were cultured in Dulbecco’s Modified Eagle Medium (DMEM) (Corning Life Sciences; Corning, NY; Cat No. 10-013-CV) supplemented with 10% v/v fetal bovine serum (FBS) (R&D Systems; Minneapolis, MN; Cat. No. S11550) and 1% v/v Penicillin-Streptomycin (Corning Life Sciences; Cat No. 10-013-CV). Cell lines were screened monthly for mycoplasma via the MycoAlertTM Mycoplasma Detection Kit (Lonza, Walkersville, MD; Cat No. LT07-318).

### Generation of BTEs and binding analysis

The BTE molecule consists of two single-chain variable regions (scFv) of antibodies, scFv145-2C11 (against murine CD3 epsilon subunit of T-cell receptor complex; CD3e)^15^ and scFv47 (against human IL13Rα2)^16, 17^, connected through a 23aa Gly_4_S flexible linker, synthesized by GenScript (Piscataway NJ) and subcloned into a pLVX-IRES-zsGreen 1 (Takara Bio, Mountain View, CA; Cat. No. 632187) lentiviral expression cassette. The CDR3 region of the scFvIL13Rα2 light chain was replaced with the CDR3 domain of a non-specific antibody MOPC-21 (designated NC-BTE) to generate a negative control protein unable to interact with IL13Rα2. For the production of soluble BTE proteins, HEK293T cells were transduced with lentiviral particles. Proteins were isolated from the supernatants using His-Tag affinity chromatography as previously described^18^, and characterized for their size and binding specificity against IL13Rα2 by Western Blot and ELISA.

### Western Blot and ELISA

Western and enzyme-linked immunosorbent assay (ELISA) were carried out as previously described^18^. Briefly, Pierce™ BCA Protein Assay Kits (ThermoFisher Scientific, Waltham, MC; Cat. No. A55864) was used to quantify the purified BTE and NC-BTE concentration. For western blot analysis, purified protein samples were then diluted into Laemmli Sample Buffer (Bio-Rad, Hercules, CA; Cat. No. 1610747) supplemented with 2.5% beta-mercaptoethanol (Sigma Aldrich, St. Louis, MO; Cat. No. M6250) and heat-denatured for 10 min at 95°C. Samples and the 250 kDa Precision Plus Protein standard (Bio-Rad Laboratories, Cat No. 1610375) were then separated by size via SDS-polyacrylamide gel electrophoresis (SDS-PAGE) using a 4-20% Midi-PROTEAN® TGX Stain-Free™ gel (Bio-Rad Laboratories; Cat. No. 17000436). The Trans-Blot® Turbo Transfer System™ (Bio-Rad Laboratories) was used to transfer separated proteins onto a methanol-activated 0.2 μm PVDF membrane (Millipore Sigma, Burlington, MA; Cat. No. IPVH08100). Membranes were blocked for 2 hours at RT with 5% non-fat dry milk in tris buffered saline-tween-20 (TBST) (Boston Bioproducts, Ashland, MA; Cat #IBB180), incubated at RT for 2 hours with HRP Anti-6X His tag® antibody (1:1000) (Abcam, Cambridge, United Kingdom; Cat. No. AD1.1.10), washed with TBST, incubated with Clarity™ Western ECL Substrate (Bio-Rad Laboratories; Cat. No. 1705061) and visualized via chemiluminescent imaging using ChemiDoc™ MP Imaging System (Bio-Rad Laboratories). For ELISA, 96 well plates were coated with 1 μg/mL of recombinant IL13Rα2hFc (R&D Systems, Minneapolis, MN, Cat No. 7147-IR-100) and IL13Rα1hFc (R&D Systems, Minneapolis, MN, Cat No. 146-IR-100). Plates were blocked with 2% bovine serum albumin in TBST, washed thrice with TBST, and incubated with various dilutions of BTE and NC-BTE. Plates were then incubated for two hours with the anti-6X His tag® antibody (1:1000) (Abcam; Cat. No. AD1.1.10), washed 5 times with TBST, and incubated with 1-Step™ Slow TMB-ELISA (Thermo Scientific; Cat. No. 34022) and 2N HCl. BTEs were quantified via a colorimetric readout at 470 nm using a BioTek plate reader (BioTek, Winooski, VT).

### Chromium-51 (Cr^51^) Release Assay

Chromium-51 (Cr^51^) release assays were used to assess BTE-induced T cell-mediated cytotoxicity against glioma cells. Briefly, splenic murine T cells were isolated from C57BL/6 mice using the EasySep™ Mouse T Cell Isolation Kit (STEMCELL Technologies; Cat. No. 19851). Glioma cells were labeled with Cr^51^ (0.05 mCi) (Revvity, Waltham, MA; Cat No. NEZ030001MC). Murine T cells were then co-cultured with labeled glioma cells at a 20:1 E:T ratio and treated with various concentrations of BTE protein. The positive control consisted of Dynabead™ CD3/CD28-activated T cells (GIBCO, Waltham, MA; Cat11452D). After 18 hours, supernatants were carefully removed, placed into 96 well LumaPlate-96 (PerkinElmer; Wellesley; MA Cat No. 6006633), dried via evaporation, and Cr^51^ activity was measured in a gamma counter (PerkinElmer; Wellesley; MA). Specific lysis was computed relative to spontaneous and maximum activity (measured via target cell incubation with 1% Triton X-100). Data was displayed as the standard error of mean of triplicates. Statistical significance was denoted based on the *p*-value from the student t-test.

### Orthotopic Xenograft and Genetically Engineered Mouse Model

All animal experiments were conducted following the Northwestern University Institutional Animal Care and Use Committee approved protocol IS00016555. Orthotopic models utilized CD45.1 and C57Bl/6 mice sourced from Jackson Laboratories. The VMDK mice were provided by Dr. J. Sampson (Duke University). The *de novo* GEMM glioma model employed nestin-Tva PTEN^fl/fl^ p53^fl/fl^ mice generously gifted by Dr. O. Becher (Mount Sinai, NY). Gliomagenesis in the GEMM was induced using DF1 cells producing RCAS-cre and by inserting PDGFB and IL13Rα2 using the RCAS-PDGFB-IL13Rα2 virus as previously described^20^. In all models, gliomas were initiated via stereotactic injection of either murine glioma cells (orthotopic model) or DF-1 viral-producing cells (GEMM) into the cerebral cortex, approximately 1 mm posterior to the bregma and 1 mm to the right of the midline, as previously described^19^. Mice were randomized by gender into treatment groups: BTE, NC-BTE, or saline. In syngeneic CT2A and GL261 models, treatments were administered intraperitoneally (i.p.) on days 7, 9, and 11 post-glioma implantations. In the de novo GEM model, treatments were given i.p. four days per week starting day 7 post-implantation for three weeks. Mice were monitored for survival with endpoints defined as either a 20% reduction in body weight or the onset of neurological symptoms.

### Flow Cytometry

Cells were blocked with Dulbecco’s Phosphate-Buffered Saline (DPBS) (Corning Life Sciences; Cat. No. 20-031-CV) supplemented with 2% FBS (R&D Systems) and TruStain FcX™ anti-mouse CD16/32 (1:200; BioLegend, San Diego, CA; Cat. No. 101319). Surface antigens were stained using fluorochrome-conjugated antibodies in 2% FBS-DPBS for 20 minutes on ice. Dead cells were labeled with eBioscience Fixable Viability Dye eFluor 780 (1:1000 in DPBS; Invitrogen, Waltham, MA; Cat. No. 65-0865-18). Cells were fixed and permeabilized with the eBioscience™ FOXP3/Transcription Factor Staining Buffer Set (Invitrogen; Cat. No. 00-5523-00), followed by intracellular staining with fluorochrome-conjugated antibodies. Data acquisition was performed using the BD FACSymphony A5-Laser Analyzer (Becton Dickinson, Franklin Lakes, NJ) at the Robert H. Lurie Comprehensive Cancer Center (RHLCCC) Flow Cytometry Core Facility. Flow cytometric analyses were conducted using FlowJo v10.9.0 software (Becton Dickinson).

For *in vitro* T cell activation experiments, splenic murine T cells were freshly isolated from C57BL/6 mice using the EasySep™ Mouse T Cell Isolation Kit (STEMCELL Technologies, Vancouver, Canada; Cat. No. 19851). Murine T cells were co-cultured with glioma cells at a 20:1 effector-to-target (E:T) ratio in triplicate wells of a 96-well plate treated with various concentrations of BTE, NC-BTE, or Dynabeads™ Mouse T-Activator CD3/CD28 (Gibco, Carlsbad, California; Cat. No. 11456D) for 24 hours. Following incubation, cells were collected for flow cytometric staining. For the TME analysis, orthotopic glioma cells were implanted and randomized into the following treatment groups: BTE, NC-BTE, or saline. Treatments were administered i.p. starting 7 days post-glioma implantation/initiation. Brains and spleens were harvested 48 hours after the final treatment and mechanically dissociated into single-cell suspensions using a 70 µm cell strainer. The cells were counted using trypan blue exclusion and processed for flow cytometric staining.

### Incucyte

For T cell activation assessments, splenic murine T cells from NOD.Nr4a1GFP/Cre mice (Jackson Laboratory, Milwaukee; RRID:IMSR JAX:037122) were used to visualize BTE-mediated T cell activation. The NOD.Nr4a1GFP/Cre mice contain a cre recombinase/green fluorescent fusion protein driven by the Nr4a1 promoter, which is a transcription factor downstream of TCR signaling. Thus, NOD.Nr4a1GFP/Cre T cells begin to express GFP following activation. Murine T cells were isolated from the mice using the EasySep™ Mouse T Cell Isolation Kit (STEMCELL Technologies; Cat. No. 19851). These murine T cells were then co-cultured with GL261-IL13Rα2 glioma cells at a 20:1 E:T ratio and treated with either BTE, NC-BTE, or Dynabeads™ Mouse T-Activator CD3/CD28. T cells were allowed to settle, and the mean fluorescent intensity at excitation of 488 nm was recorded and quantified from 12 images obtained between 0.75 and 18 hours post-treatment.

### Histology

The protocol for detecting immune cells in the murine brain of the GEMM has been previously described^20^. The following antibodies were used: IL13Rα2 (R&D Systems, Cat No. AF146), Olig2 (Abcam, ab109186), Ki67 (Abcam, ab16667), CD3 (Abcam, ab16669), and CD11b (Abcam, ab133357). For the detection of CD8^+^ T cells in the brains of long-term surviving mice, 4% paraformaldehyde (PFA)-fixed paraffin-embedded 4 μm thick-tissue section from the mouse brains were incubated with the CD8a monoclonal antibody (4SM16) (Invitrogen, 14-0195-82), followed by 3,3′-diaminobenzidine chromogen staining. The quantification of the stained areas was performed using the ImageJ software. The tissue was stained with hematoxylin and eosin (H&E) to highlight the morphological changes, and the image analysis was performed using the NDP.view 2.8.24 software (Hamamatsu Photonics K.K.).

### Magnetic Resonance Imaging

MRI of glioma-bearing mice was conducted using a Bruker Clinscan 7T system equipped with a dedicated multi-channel brain coil for high sensitivity and spatial resolution of whole-brain images. Anesthesia was delivered via a built-in MRI holder nose cone (100% oxygen mixed with isoflurane). Appropriate physiological conditions (i.e., body temperature and respiration rate) were maintained throughout each scanning session using the dedicated monitoring and control systems (Sai Instruments, NJ). Advanced Siemens MR sequences (Syngo VB15 Imaging platform, Siemens) were used to obtain anatomical whole-brain 3D T1-weighted images with ∼ 100-micron isotropic spatial resolution obtained using a FLASH-based 3D gradient-echo sequence with TR=100 msec and TE=3 msec. 2D quantitative multiple echo-time spin-echo sequences (TR= 1500 msec and 12 echo times TE ranging from 10-120 msec) were used to acquire whole-brain images. Following the acquisition, DICOM images were exported to Northwestern Feinberg Medical School cloud-based repository (FSMresFiles Server) and processed using a combination of shareware and commercial imaging software (Jim7.0 Xinapse, ITK_SNAP). T2 maps were extracted by fitting multiple sets of images with different echo times on a voxel-by-voxel bases using a least-square fitting tool included in JIM7.0 Xinapse image processing software was used with the expectation of using the regional quantitative variability of the extracted maps as a marker of intratumor BiTE-induced effects. A regional semi-automated threshold-based segmentation tool included in the ITK-SNAP software was used to extract 3D whole-tumor volumetrics. Tumor quantitative T2 parametric maps were normalized using T2 values from normal (i.e., no tumor) brain tissue. The resulting T2tumor/T2tissue normalized parametric maps were then analyzed using a threshold-based approach, which defined quantitative threshold windows (from high vs low T2tumor/T2tissue values). The resulting segmented regions corresponding to different ranges of T2tumor/T2tissue values were color-coded, providing a spatial representation of TME regions with different biological properties. This approach enabled the identification and use of a robust non-invasive imaging metric that has the potential when compared with the corresponding histopathological analysis conducted ex vivo on the same tumors, to facilitate interpretation of the different tumor subregions (i.e., different tumor habitats), which are reflective of different stages of progression/viability. The volume of each sub-region (i.e., regions with specific average T2tumor/T2tissue values) for each mouse was used to generate quantitative dimensional parameters (using whole brain volume as a normalization factor) that were used for statistical analysis. Computed and compared average indexes for the control and the BTE-treated groups were made for the high and low T2 ranges.

Based on existing preclinical and clinical reports suggesting the predictive diagnostic potential of regional intra-tumor T2 values, we extracted and calculated a quantitative dimensional parameter (i.e. volumetric ratios) that could potentially be used as a non-invasive biomarker of tumor progression and/or response to therapy. This parameter was defined as the Tumor Therapeutic Response Index and was calculated as follows: *Volume Low Range T2 tumor/T2tissue/Volume Whole Brain*. The driving hypothesis is that lower tumor tissue T2 values tend to be detected (clinically and pre-clinically) in less active or therapeutically more responsive tumors, thus leading to consistently high Tumor Therapeutic Response Index values in tumors of the BTE-treated mice compared to the control tumors.

### Single Cell RNA Sequencing

Single-cell RNA sequencing (scRNA-seq) was performed on both control and BTE-treated gliomas from the Nestin-Tva-PTEN^fl/fl^-P53^fl/fl^ mice from two independent experiments with identical study design. These mice were randomized evenly by gender into two groups, with 5 mice per group. Seven days post glioma induction, mice were treated with either saline or BTE (10mg/kg) via i.p. route four times per week for 3 weeks. Forty-eight hours after the last treatment, mice were anesthetized and perfused intracardially with 10 ml of saline. The glioma tissue was excised and placed into single-cell suspension using the Adult Brain Dissociation Kit, Mouse and Rat (Miltenyi Biotec, Gaithersburg, MD; Cat No. 130-107-677). Single-cell suspensions were then incubated with anti-CD16/32 (Fc Block) antibodies (BioLegend) and then subjected to anti-mouse CD45 bead-based separation (Miltenyi Biotec; Cat. No 130-052-301). The scRNA-seq was performed on the glioma-infiltrating immune cells enriched at a CD45+ to CD45-cell ratio of 8:2 at a 4000 live cells/µL concentration.

The PIPseq™ T2 3’ Single Cell RNA Kit v4.0 kit (Fluent Biosciences, Cat No. FBS-SCR-T20-4-V4.05) was used for scRNA-seq prep. PIPseq uses a microfluidics-free, templated emulsification technology to generate monodispersed cell droplets. 4×10^4^ cells per sample were added to particle-templated instant partitions, emulsified via rapid mechanical vertexing, lysed to capture mRNA on barcoded poly(T) decorated beads, transcribed into cDNA, amplified with PCR, and sample indexed with unique P7 and P5 Illumina compatible indexes. The barcoded cDNA libraries were then subjected to pooled sequencing on an Illumina NovaSeq X Plus platform at a depth of 650 million reads per sample at Novogene (Sacramento, CA). Demultiplexed FASTq files were aligned to the Human Hg38 genome assembly (GRCh38.p13, 2022) using the PIPseeker (Fluent BioSciences) software. The resulting data matrix files were analyzed for each sample in R (v4.2.3) using the Seurat (v5) package. Cells were filtered with >2.5% mitochondrial read counts with >500 expressed genes. The scDblFinder (v1.18.0) was used to identify and remove doublets. In Seurat, the data was log normalized using a scale factor of 10,000. Default parameters were used to identify highly variable genes. Principal component analysis was conducted on the top 2000 variable genes. The top 17 principal components were selected for neighboring and clustering identification. To correct the batch effect between the two experiments, canonical correlation analysis (CCA) was performed. Subsequently the JoinLayers function was employed to merge sample layers. The clusters were visualized in Uniform Manifold Approximation and Projection (UMAP) of the first 17 CCA reductions. FindAllMarkers was used to find top differentially expressed genes between clusters and identify cell types based on canonical markers. Seurat workflow was used to subcluster immune cells and the FindMarkers function was used to identify differentially expressed genes (DEGs) between treatment groups. The ClusterProfiler (v4.12.0) was used to perform gene ontology enrichment analysis on significant differentially expressed genes DEGs. The R package ggplot2 (v3.5.1) and ComplexHeatmap (v2.20.0) was used to construct the plots and heat maps.

### Automated multiplexed sequential immunofluorescence imaging

Automated hyperplex immunofluorescence staining and imaging was performed on FFPE sections using the COMET™ system (Lunaphore Technologies). The following targets and the associated antibodies were used for this study: GFAP (Sigma, Clone GA5, MAB360, 1/3000 dilution); CD31 (Abcam, Clone EPR17259, AB225883, 1/1500 dilution); CD4 (Abcam, Clone EPR19514, Ab183685, 1/500 dilution; CD8 (Cell Signaling, Clone D4W2Z XP, 98941, dilution 1/500); IFN-γ (Bioss, BS-0480R, polyclonal, dilution 1/100); TCF1 (Cell signaling, clone C63D9, cat#2203, dilution 1/500); IL13Rα2 (Cell Signaling, E7U7B, 85677, dilution 1/200). All antibodies were validated using conventional immunohistochemistry or IF staining in conjunction with corresponding fluorophores and 4’,6-diamidino-2-pheynlindole counterstain (DAPI, ThermoFisher Scientific). For optimal concentration and best signal-to-noise ratio, all antibodies were tested at 3 different dilutions, starting with the manufacturer-recommended dilution (MRD), MRD/2, and MRD/4. Secondary Alexa fluorophore 555 (ThermoFisher Scientific) and Alexa fluorophore 647 (ThermoFisher Scientific) were used at 1/200 and 1/400 dilutions, respectively. The optimizations and full runs of the multiplexed panel were executed using the Lunaphore COMET™ platform (characterization 2 and 3 protocols, and seqIF™ protocols, respectively). The workflow was performed in parallel with automated iterative cycles of staining, followed by imaging, and elution of the primary and secondary antibodies. No sample manipulation is required during the entire workflow. All reagents were diluted in Multistaining Buffer (BU06, Lunaphore Technologies). The Elution step lasted 2 minutes for each cycle and was performed with Elution Buffer (BU07-L, Lunaphore Technologies) at 37°C. The Quenching step lasted for 30 seconds and was performed with Quenching Buffer (BU08-L, Lunaphore Technologies). The Imaging step was performed with Imaging Buffer (BU09, Lunaphore Technologies) with exposure times set at 4 minutes for all primary antibodies and secondary antibodies at 2 minutes. The Imaging step is performed with an integrated epifluorescent microscope at 20x magnification. Image registration was performed immediately after concluding the IF staining and imaging procedures by COMET™ Control Software. Each seqIF™ protocol resulted in a multi-stack OME-TIFF file where the imaging outputs from each cycle are stitched and aligned. COMET™ OME-TIFF file contains DAPI image, intrinsic tissue autofluorescence in the TRITC and Cy5 channels, and a single fluorescent layer per marker. Markers were subsequently pseudo-colored for visualization of multiplexed antibodies using HORIZON Viewer™ software.

### Statistical analyses

All statistical analysis was completed in R (v4.2.3). Statistical analysis was performed by one-way ANOVA and p values were adjusted for multiple comparisons using Tukey’s test. The Kaplan-Meier method was used to compare survival between treatment groups and analyzed with the log-rank (Mantle-Cox) test. p values were adjusted for multiple comparisons by Bonferroni’s correction.

## Acknowledgments

This study was partly supported by the following grants: NINDS R33 NS101150, R01 NS106379, and R01 NS122395 (IVB). The authors thank the Mouse Histology and Phenotyping Laboratory and the Small Animal Imaging Cores supported by National Cancer Institute Grant P30-CA060553, awarded to the Robert H. Lurie Comprehensive Cancer Center.

## Author contributions

IVB conceived the study and provided funding; MZ, JTD, IVB designed experiments; MZ, JTD, DP, HN and RNL performed experiments; DH and BZ provided expertise on GEM and Nur77 mouse models; MZ, JTD, DP, HN, LL, C.L-C, JM, and IVB performed data analysis; JTD, DP, and IVB co-wrote the manuscript. JTD, AH, IVB performed revision of the manuscript. MZ and JTD contributed equally to this work. All authors met the criteria for authorship and approved the final manuscript for publication.

## References

1. Alexander BM, Cloughesy TF. Adult glioblastoma. Journal of Clinical Oncology. 2017;35(21):2402–9.

2. Stupp R, Mason WP, Van Den Bent MJ, Weller M, Fisher B, Taphoorn MJ, Belanger K, Brandes AA, Marosi C, Bogdahn U. Radiotherapy plus concomitant and adjuvant temozolomide for glioblastoma. New England journal of medicine. 2005;352(10):987–96.

3. Bi WL, Beroukhim R. Beating the odds: extreme long-term survival with glioblastoma. Society for Neuro-Oncology; 2014. p. 1159–60.

4. Stupp R, Taillibert S, Kanner A, Read W, Steinberg DM, Lhermitte B, Toms S, Idbaih A, Ahluwalia MS, Fink K. Effect of tumor-treating fields plus maintenance temozolomide vs maintenance temozolomide alone on survival in patients with glioblastoma: a randomized clinical trial. Jama. 2017;318(23):2306–16.

5. Alsajjan R, Mason WP. Bispecific T-cell engagers and chimeric antigen receptor T-cell therapies in glioblastoma: an update. Current Oncology. 2023;30(9):8501–49.

6. Gedeon PC, Schaller TH, Chitneni SK, Choi BD, Kuan CT, Suryadevara CM, Snyder DJ, Schmittling RJ, Szafranski SE, Cui X, Healy PN, Herndon JE, 2nd, McLendon RE, Keir ST, Archer GE, Reap EA, Sanchez-Perez L, Bigner DD, Sampson JH. A Rationally Designed Fully Human EGFRvIII:CD3-Targeted Bispecific Antibody Redirects Human T Cells to Treat Patient-derived Intracerebral Malignant Glioma. Clin Cancer Res. 2018;24(15):3611–31. Epub 20180427. doi: 10.1158/1078-0432.CCR-17-0126. PubMed PMID: 29703821; PMCID: PMC6103776.

7. Park DH, Bhojnagarwala PS, Liaw K, Bordoloi D, Tursi NJ, Zhao S, Binder ZA, O’Rourke D, Weiner DB. Novel tri-specific T-cell engager targeting IL-13Rα2 and EGFRvIII provides long-term survival in heterogeneous GBM challenge and promotes antitumor cytotoxicity with patient immune cells. Journal for ImmunoTherapy of Cancer. 2024;12(12):e009604. doi: 10.1136/jitc-2024-009604.

8. Park DH, Liaw K, Bhojnagarwala P, Zhu X, Choi J, Ali AR, Bordoloi D, Gary EN, O’Connell RP, Kulkarni A. Multivalent in vivo delivery of DNA-encoded bispecific T cell engagers effectively controls heterogeneous GBM tumors and mitigates immune escape. Molecular Therapy-Oncolytics. 2023;28:249–63.

9. Gedeon PC, Streicker MA, Schaller TH, Archer GE, Jokinen MP, Sampson JH. GLP toxicology study of a fully-human T cell redirecting CD3: EGFRvIII binding immunotherapeutic bispecific antibody. Plos one. 2020;15(7):e0236374.

10. Sonabend AM, Gould A, Amidei C, Ward R, Schmidt KA, Zhang DY, Gomez C, Bebawy JF, Liu BP, Bouchoux G, Desseaux C, Helenowski IB, Lukas RV, Dixit K, Kumthekar P, Arrieta VA, Lesniak MS, Carpentier A, Zhang H, Muzzio M, Canney M, Stupp R. Repeated blood-brain barrier opening with an implantable ultrasound device for delivery of albumin-bound paclitaxel in patients with recurrent glioblastoma: a phase 1 trial. Lancet Oncol. 2023;24(5):509–22. doi: 10.1016/S1470-2045(23)00112-2. PubMed PMID: 37142373; PMCID: PMC10256454.

11. Yin Y, Rodriguez JL, Li N, Thokala R, Nasrallah MP, Hu L, Zhang L, Zhang JV, Logun MT, Kainth D, Haddad L, Zhao Y, Wu T, Johns EX, Long Y, Liang H, Qi J, Zhang X, Binder ZA, Lin Z, O’Rourke DM. Locally secreted BiTEs complement CAR T cells by enhancing killing of antigen heterogeneous solid tumors. Mol Ther. 2022;30(7):2537–53. Epub 20220514. doi: 10.1016/j.ymthe.2022.05.011. PubMed PMID: 35570396; PMCID: PMC9263323.

12. Choi BD, Yu X, Castano AP, Bouffard AA, Schmidts A, Larson RC, Bailey SR, Boroughs AC, Frigault MJ, Leick MB. CAR-T cells secreting BiTEs circumvent antigen escape without detectable toxicity. Nature biotechnology. 2019;37(9):1049–58.

13. Bagley SJ, Logun M, Fraietta JA, Wang X, Desai AS, Bagley LJ, Nabavizadeh A, Jarocha D, Martins R, Maloney E. Intrathecal bivalent CAR T cells targeting EGFR and IL13Rα2 in recurrent glioblastoma: phase 1 trial interim results. Nature Medicine. 2024;30(5):1320–9.

14. Choi BD, Gerstner ER, Frigault MJ, Leick MB, Mount CW, Balaj L, Nikiforow S, Carter BS, Curry WT, Gallagher K. Intraventricular CARv3-TEAM-E T cells in recurrent glioblastoma. New England Journal of Medicine. 2024;390(14):1290–8.

15. Leo O, Foo M, Sachs DH, Samelson LE, Bluestone JA. Identification of a monoclonal antibody specific for a murine T3 polypeptide. Proc Natl Acad Sci U S A. 1987;84(5):1374–8. doi: 10.1073/pnas.84.5.1374. PubMed PMID: 2950524; PMCID: PMC304432.

16. Balyasnikova IV, Wainwright DA, Solomaha E, Lee G, Han Y, Thaci B, Lesniak MS. Characterization and immunotherapeutic implications for a novel antibody targeting interleukin (IL)-13 receptor α2. J Biol Chem. 2012;287(36):30215–27. Epub 20120709. doi: 10.1074/jbc.M112.370015. PubMed PMID: 22778273; PMCID: PMC3436275.

17. Kim JW, Young JS, Solomaha E, Kanojia D, Lesniak MS, Balyasnikova IV. A novel single-chain antibody redirects adenovirus to IL13Rα2-expressing brain tumors. Sci Rep. 2015;5:18133. Epub 20151214. doi: 10.1038/srep18133. PubMed PMID: 26656559; PMCID: PMC4677343.

18. Pituch KC, Zannikou M, Ilut L, Xiao T, Chastkofsky M, Sukhanova M, Bertolino N, Procissi D, Amidei C, Horbinski CM, Aboody KS, James CD, Lesniak MS, Balyasnikova IV. Neural stem cells secreting bispecific T cell engager to induce selective antiglioma activity. Proc Natl Acad Sci U S A. 2021;118(9). Epub 2021/02/26. doi: 10.1073/pnas.2015800118. PubMed PMID: 33627401; PMCID: PMC7936285.

19. Pituch KC, Miska J, Krenciute G, Panek WK, Li G, Rodriguez-Cruz T, Wu M, Han Y, Lesniak MS, Gottschalk S, Balyasnikova IV. Adoptive Transfer of IL13Ralpha2-Specific Chimeric Antigen Receptor T Cells Creates a Pro-inflammatory Environment in Glioblastoma. Mol Ther. 2018;26(4):986–95. Epub 20180208. doi: 10.1016/j.ymthe.2018.02.001. PubMed PMID: 29503195; PMCID: PMC6079480.

20. Seblani M, Zannikou M, Duffy JT, Levine RN, Liu Q, Horbinski CM, Becher OJ, Balyasnikova IV. A New Mouse Model of Diffuse Midline Glioma to Test Targeted Immunotherapies. bioRxiv. 2021:2021.10.15.464284. doi: 10.1101/2021.10.15.464284.

21. Dreier T, Lorenczewski G, Brandl C, Hoffmann P, Syring U, Hanakam F, Kufer P, Riethmuller G, Bargou R, Baeuerle PA. Extremely potent, rapid and costimulation-independent cytotoxic T-cell response against lymphoma cells catalyzed by a single-chain bispecific antibody. International journal of cancer. 2002;100(6):690–7.

22. Offner S, Hofmeister R, Romaniuk A, Kufer P, Baeuerle PA. Induction of regular cytolytic T cell synapses by bispecific single-chain antibody constructs on MHC class I-negative tumor cells. Molecular immunology. 2006;43(6):763–71.

23. Przepiorka D, Ko C-W, Deisseroth A, Yancey CL, Candau-Chacon R, Chiu H-J, Gehrke BJ, Gomez-Broughton C, Kane RC, Kirshner S. FDA approval: blinatumomab. Clinical Cancer Research. 2015;21(18):4035–9.

24. Liu J, Zhu J. Progresses of T-cell-engaging bispecific antibodies in treatment of solid tumors. International Immunopharmacology. 2024;138:112609.

25. Dhillon S. Tarlatamab: First Approval. Drugs. 2024:1–9.

26. Dhillon S. Tebentafusp: first approval. Drugs. 2022;82(6):703–10.

27. Quail DF, Joyce JA. The microenvironmental landscape of brain tumors. Cancer cell. 2017;31(3):326–41.

28. Bowman RL, Klemm F, Akkari L, Pyonteck SM, Sevenich L, Quail DF, Dhara S, Simpson K, Gardner EE, Iacobuzio-Donahue CA. Macrophage ontogeny underlies differences in tumor-specific education in brain malignancies. Cell reports. 2016;17(9):2445–59.

29. Wang G, Zhong K, Wang Z, Zhang Z, Tang X, Tong A, Zhou L. Tumor-associated microglia and macrophages in glioblastoma: From basic insights to therapeutic opportunities. Frontiers in immunology. 2022;13:964898.

30. Brown CE, Alizadeh D, Starr R, Weng L, Wagner JR, Naranjo A, Ostberg JR, Blanchard MS, Kilpatrick J, Simpson J. Regression of glioblastoma after chimeric antigen receptor T-cell therapy. New England Journal of Medicine. 2016;375(26):2561–9.

31. Krenciute G, Prinzing BL, Yi Z, Wu M-F, Liu H, Dotti G, Balyasnikova IV, Gottschalk S. Transgenic expression of IL15 improves antiglioma activity of IL13Rα2-CAR T cells but results in antigen loss variants. Cancer immunology research. 2017;5(7):571–81.

32. van de Donk NW, Zweegman S. T-cell-engaging bispecific antibodies in cancer. The Lancet. 2023;402(10396):142–58.

33. Cattaruzza F, Nazeer A, To M, Hammond M, Koski C, Liu LY, Pete Yeung V, Rennerfeldt DA, Henkensiefken A, Fox M. Precision-activated T-cell engagers targeting HER2 or EGFR and CD3 mitigate on-target, off-tumor toxicity for immunotherapy in solid tumors. Nature Cancer. 2023;4(4):485–501.

34. Zeng J, Zhang J, Yang Y-Z, Wang F, Jiang H, Chen H-D, Wu H-Y, Sai K, Hu W-M. IL13RA2 is overexpressed in malignant gliomas and related to clinical outcome of patients. American journal of translational research. 2020;12(8):4702.

35. Joshi BH, Plautz GE, Puri RK. Interleukin-13 receptor α chain: a novel tumor-associated transmembrane protein in primary explants of human malignant gliomas. Cancer research. 2000;60(5):1168–72.

36. Bhardwaj R, Suzuki A, Leland P, Joshi BH, Puri RK. Identification of a novel role of IL-13Rα2 in human Glioblastoma multiforme: interleukin-13 mediates signal transduction through AP-1 pathway. Journal of Translational Medicine. 2018;16:1–13.

37. Kawakami K, Leland P, Puri RK. Structure, function, and targeting of interleukin 4 receptors on human head and neck cancer cells. Cancer research. 2000;60(11):2981–7.

38. Kawakami M, Kawakami K, Kasperbauer JL, Hinkley LL, Tsukuda M, Strome SE, Puri RK. Interleukin-13 receptor α2 chain in human head and neck cancer serves as a unique diagnostic marker. Clinical cancer research. 2003;9(17):6381–8.

39. Bartolomé RA, Martín-Regalado Á, Jaén M, Zannikou M, Zhang P, de Los Ríos V, Balyasnikova IV, Casal JI. Protein tyrosine phosphatase-1B inhibition disrupts IL13Rα2-promoted invasion and metastasis in cancer cells. Cancers. 2020;12(2):500.

40. Fujisawa T, Joshi B, Nakajima A, Puri RK. A novel role of interleukin-13 receptor α2 in pancreatic cancer invasion and metastasis. Cancer research. 2009;69(22):8678–85.

41. Okamoto H, Yoshimatsu Y, Tomizawa T, Kunita A, Takayama R, Morikawa T, Komura D, Takahashi K, Oshima T, Sato M. Interleukin-13 receptor α2 is a novel marker and potential therapeutic target for human melanoma. Scientific reports. 2019;9(1):1281.

42. Papageorgis P, Ozturk S, Lambert AW, Neophytou CM, Tzatsos A, Wong CK, Thiagalingam S, Constantinou AI. Targeting IL13Ralpha2 activates STAT6-TP63 pathway to suppress breast cancer lung metastasis. Breast Cancer Research. 2015;17:1–15.

43. Márquez-Ortiz RA, Contreras-Zárate MJ, Tesic V, Alvarez-Eraso KL, Kwak G, Littrell Z, Costello JC, Sreekanth V, Ormond DR, Karam SD. IL13Rα2 promotes proliferation and outgrowth of breast cancer brain metastases. Clinical Cancer Research. 2021;27(22):6209–21.

44. Shikalov A, Koman I, Kogan NM. Targeted glioma therapy—clinical trials and future directions. Pharmaceutics. 2024;16(1):100.

45. Bhojnagarwala PS, O’Connell RP, Park D, Liaw K, Ali AR, Bordoloi D, Cassel J, Tursi NJ, Gary E, Weiner DB. In vivo DNA-launched bispecific T cell engager targeting IL-13Ralpha2 controls tumor growth in an animal model of glioblastoma multiforme. Mol Ther Oncolytics. 2022;26:289–301. Epub 20220706. doi: 10.1016/j.omto.2022.07.003. PubMed PMID: 36090479; PMCID: PMC9418050.

46. O’Rourke DM, Nasrallah MP, Desai A, Melenhorst JJ, Mansfield K, Morrissette JJ, Martinez-Lage M, Brem S, Maloney E, Shen A. A single dose of peripherally infused EGFRvIII-directed CAR T cells mediates antigen loss and induces adaptive resistance in patients with recurrent glioblastoma. Science translational medicine. 2017;9(399):eaaa0984.

47. Escobar G, Mangani D, Anderson AC. T cell factor 1: A master regulator of the T cell response in disease. Science immunology. 2020;5(53):eabb9726.

48. Okła K, Farber DL, Zou W. Immune Memory Focus: Tissue-resident memory T cells in tumor immunity and immunotherapy. The Journal of experimental medicine. 2021;218(4).

49. Crane AT, Chrostek MR, Krishna VD, Shiao M, Toman NG, Pearce CM, Tran SK, Sipe CJ, Guo W, Voth JP. Zika virus-based immunotherapy enhances long-term survival of rodents with brain tumors through upregulation of memory T-cells. PLoS One. 2020;15(10):e0232858.

50. Ning J, Gavil NV, Wu S, Wijeyesinghe S, Weyu E, Ma J, Li M, Grigore F-N, Dhawan S, Skorput AG. Functional virus-specific memory T cells survey glioblastoma. Cancer Immunology, Immunotherapy. 2022:1–13.

51. Zhao Q, Hu J, Kong L, Jiang S, Tian X, Wang J, Hashizume R, Jia Z, Fowlkes NW, Yan J. FGL2-targeting T cells exhibit antitumor effects on glioblastoma and recruit tumor-specific brain-resident memory T cells. Nature communications. 2023;14(1):735.

52. La Manna MP, Di Liberto D, Lo Pizzo M, Mohammadnezhad L, Shekarkar Azgomi M, Salamone V, Cancila V, Vacca D, Dieli C, Maugeri R. The abundance of tumor-infiltrating CD8+ tissue resident memory T lymphocytes correlates with patient survival in glioblastoma. Biomedicines. 2022;10(10):2454.

53. Corgnac S, Malenica I, Mezquita L, Auclin E, Voilin E, Kacher J, Halse H, Grynszpan L, Signolle N, Dayris T. CD103+ CD8+ TRM cells accumulate in tumors of anti-PD-1-responder lung cancer patients and are tumor-reactive lymphocytes enriched with Tc17. Cell Reports Medicine. 2020;1(7).

54. Edwards J, Wilmott JS, Madore J, Gide TN, Quek C, Tasker A, Ferguson A, Chen J, Hewavisenti R, Hersey P. CD103+ tumor-resident CD8+ T cells are associated with improved survival in immunotherapy-naïve melanoma patients and expand significantly during anti–PD-1 treatment. Clinical Cancer Research. 2018;24(13):3036–45.

55. Wang M, Pruteanu I, Cohen AD, Garfall AL, Milone MC, Tian L, Gonzalez VE, Gill S, Frey NV, Barrett DM. Identification and validation of predictive biomarkers to CD19-and BCMA-specific CAR T-cell responses in CAR T-cell precursors. Blood. 2019;134:622.

56. Mathios D, Srivastava S, Kim T, Bettegowda C, Lim M. Emerging technologies for non-invasive monitoring of treatment response to immunotherapy for brain tumors. Neuromolecular medicine. 2022;24(2):74–87.

57. Sinigaglia M, Assi T, Besson FL, Ammari S, Edjlali M, Feltus W, Rozenblum-Beddok L, Zhao B, Schwartz LH, Mokrane F-Z. Imaging-guided precision medicine in glioblastoma patients treated with immune checkpoint modulators: research trend and future directions in the field of imaging biomarkers and artificial intelligence. EJNMMI research. 2019;9:1–20.

58. Xie T, Chen X, Fang J, Xue W, Zhang J, Tong H, Liu H, Guo Y, Yang Y, Zhang W. Non-invasive monitoring of the kinetic infiltration and therapeutic efficacy of nanoparticle-labeled chimeric antigen receptor T cells in glioblastoma via 7.0-Tesla magnetic resonance imaging. Cytotherapy. 2021;23(3):211–22.

59. Booth TC, Larkin TJ, Yuan Y, Kettunen MI, Dawson SN, Scoffings D, Canuto HC, Vowler SL, Kirschenlohr H, Hobson MP. Analysis of heterogeneity in T2-weighted MR images can differentiate pseudoprogression from progression in glioblastoma. PLoS One. 2017;12(5):e0176528.

60. Radbruch A, Lutz K, Wiestler B, Bäumer P, Heiland S, Wick W, Bendszus M. Relevance of T2 signal changes in the assessment of progression of glioblastoma according to the Response Assessment in Neurooncology criteria. Neuro-oncology. 2011;14(2):222–9.

61. Reuter G, Lommers E, Balteau E, Simon J, Phillips C, Scholtes F, Martin D, Lombard A, Maquet P. Multiparameter quantitative histological MRI values in high-grade gliomas: a potential biomarker of tumor progression. Neuro-Oncology Practice. 2020;7(6):646–55.

62. Mestas J, Hughes CC. Of mice and not men: differences between mouse and human immunology. The Journal of Immunology. 2004;172(5):2731–8.

63. Ueda O, Wada NA, Kinoshita Y, Hino H, Kakefuda M, Ito T, Fujii E, Noguchi M, Sato K, Morita M. Entire CD3ε, δ, and γ humanized mouse to evaluate human CD3–mediated therapeutics. Scientific reports. 2017;7(1):45839.

64. Miescher GC, Schreyer M, MacDonald HR. Production and characterization of a rat monoclonal antibody against the murine CD3 molecular complex. Immunology letters. 1989;23(2):113–8.

65. Schaller TH, Foster MW, Thompson JW, Spasojevic I, Normantaite D, Moseley MA, Sanchez-Perez L, Sampson JH. Pharmacokinetic analysis of a novel human EGFRvIII: CD3 bispecific antibody in plasma and whole blood using a high-resolution targeted mass spectrometry approach. Journal of proteome research. 2019;18(8):3032–41.

66. Suurs FV, Lorenczewski G, Bailis JM, Stienen S, Friedrich M, Lee F, van der Vegt B, de Vries EG, de Groot DJA, Lub-de Hooge MN. Mesothelin/CD3 Half-Life–Extended Bispecific T-Cell Engager Molecule Shows Specific Tumor Uptake and Distributes to Mesothelin and CD3-Expressing Tissues. Journal of Nuclear Medicine. 2021;62(12):1797–804.

67. Lorenczewski G, Friedrich M, Kischel R, Dahlhoff C, Anlahr J, Balazs M, Rock D, Boyle MC, Goldstein R, Coxon A. Generation of a half-life extended anti-CD19 BiTE® antibody construct compatible with once-weekly dosing for treatment of CD19-positive malignancies. Blood. 2017;130:2815.

68. Arvedson TL, Balazs M, Bogner P, Black K, Graham K, Henn A, Friedrich M, Hoffmann P, Kischel R, Kufer P. Generation of half-life extended anti-CD33 BiTE® antibody constructs compatible with once-weekly dosing. Cancer Research. 2017;77(13_Supplement):55-.

69. Mandrup OA, Ong SC, Lykkemark S, Dinesen A, Rudnik-Jansen I, Dagnæs-Hansen NF, Andersen JT, Alvarez-Vallina L, Howard KA. Programmable half-life and anti-tumour effects of bispecific T-cell engager-albumin fusions with tuned FcRn affinity. Communications biology. 2021;4(1):310.

