## Supplemental Figures for "A Bi-Specific T Cell-Engaging Antibody Triggers Protective Immune Memory and Glioma Microenvironment Remodeling in Immune Competent Preclinical Models"

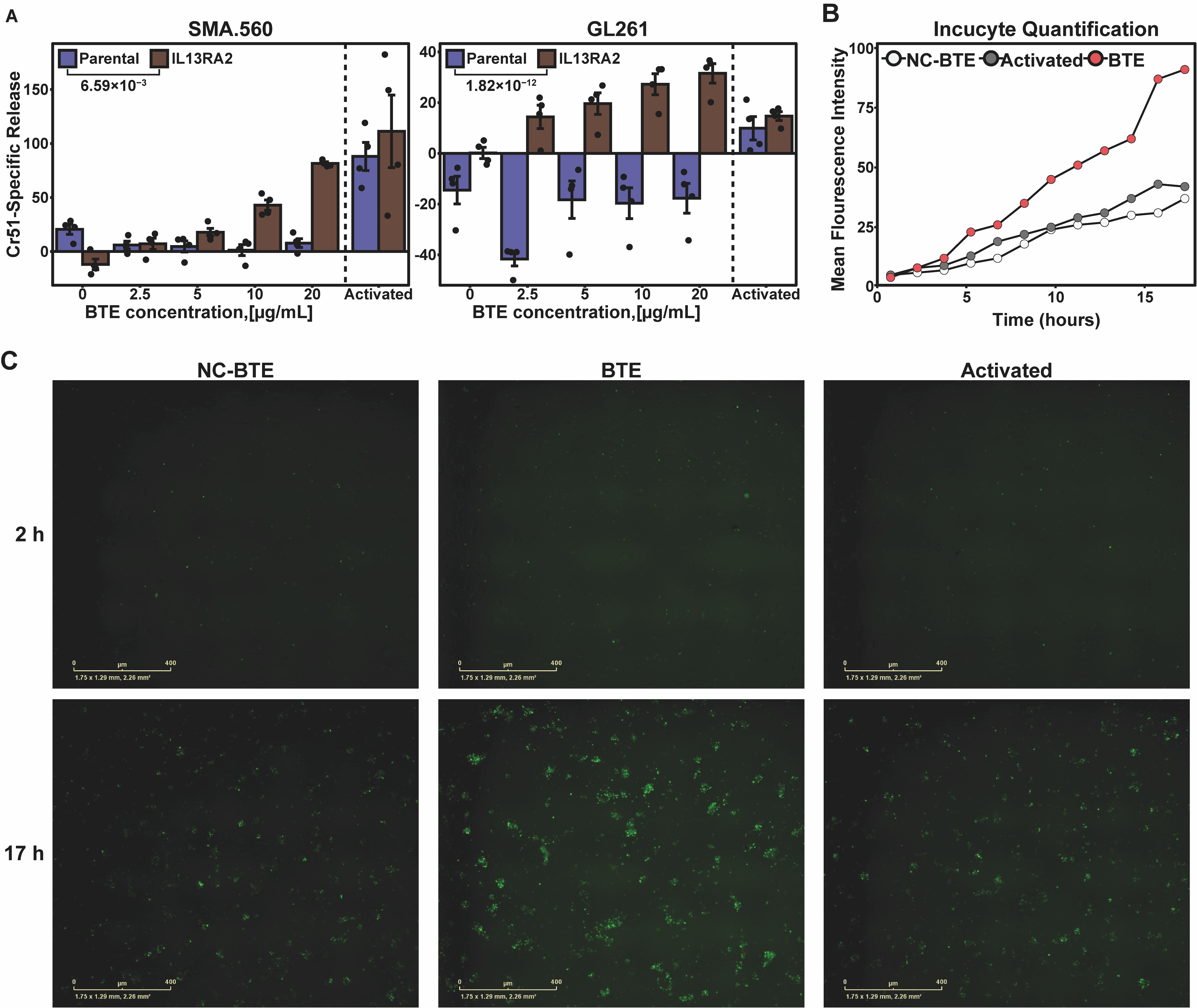

**Supplementary Figure 1. Generation and Functional Characterization of CD3xIL13R**α**2 BTE.** (*A*) Cr51 release assay of Chromium-51 (Cr51) release assay of either parental (blue) or IL13Rα2 modified (brown) SMA560 (left) and GL261(right) murine glioma cells co-cultured with murine T cells and treated with varying doses of BTE or Dynabeads™ Mouse T-Activator CD3/CD28 (Activated) showing BTE mediated killing is IL13Rα2 antigen-dependent (Effector-to-target [E: T] ratio = 20:1; Incubation time = 24 hr; n=3/group). (*B*) Quantification of mean fluorescent intensities from Incucyte NOD.Nr4a1GFP/Cre T cell activation experiment. (*C*) Representative Incucyte-generated images of murine T cells isolated from NOD.Nr4a1GFP/Cre mice cocultured with GL261-Il13Rα2 glioma cells at 2 hours and 17 hours post-treatment with NC-BTE (left), BTE (middle), and Dynabeads™ Mouse T-Activator CD3/CD28 (right). Briefly, Nr4a1 is a transcription factor that is downstream of TCR signaling. NOD.Nr4a1GFP/Cre mice contain a cre recombinase/green fluorescent fusion protein driven by the Nr4a1 (e.g., Nur77) promoter. Thus, NOD.Nr4a1GFP/Cre T cells began to express GFP following activation and were used as reporter assays to visualize BTE-mediated T cell activation.

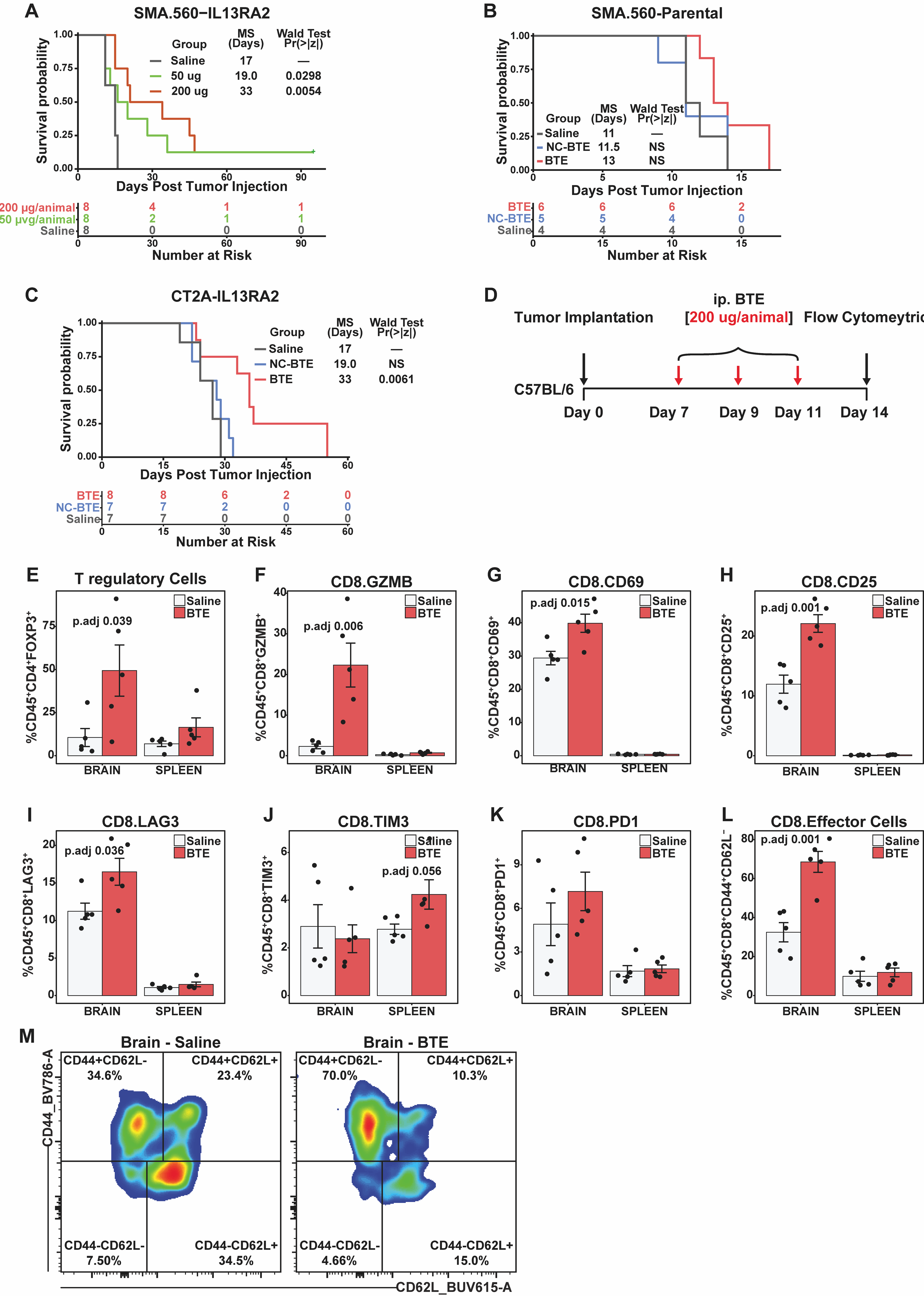

**Supplementary Figure 2. BTE Enhances Survival and Modulates Tumor Microenvironment of Mice Bearing IL13R**α**2-expressing murine glioma.** (*A*) Kaplan–Meier survival curve of SMA.560-IL13Rα2 tumor-bearing mice treated with BTE at 200 µg/animal (red), BTE (50 µg/animal) (green), or saline (black). Wald test from Cox proportional hazards regression analysis. (*B*) Kaplan–Meier survival curve of parental SMA.560 tumor-bearing mice treated with BTE at 200 µg/animal (red), NC-BTE (200 µg/animal) (blue), or saline (black). Wald test from Cox proportional hazards regression analysis (*C*) Kaplan–Meier survival curve of CT2A-IL13Rα2 tumor-bearing mice treated with BTE at 200 µg/animal (red), NC-BTE (200 µg/animal) (blue), or saline (black). Wald test from Cox proportional hazards regression analysis. (*D*) An experimental setup was used in flow cytometric analysis of the brains of SMA.560-IL13Rα2 tumor-bearing mice following treatment with either BTE or NC-BTE (n=5/group). Flow cytometric analysis revealed (*E*) increase in regulatory T cell (CD45+CD4+FOXP3+), (*F*) increase in cytotoxic CD8 T cell (CD45+CD8+GZMB+), (*G*) increase in activated CD69 expressing (CD45+CD8+CD69+) and (*H*) activated CD25 expressing CD8 T cells (CD45+CD8+CD25+), (*I*) increase in Lag3 expressing CD8 T cells (CD45+CD8+ Lag3+), (*J*) no change in Tim3 expressing CD8 T cells (CD45+ CD8+ Tim3+), (*K*) no change in Pdcd1 expressing CD8 T cells (CD45+ CD8+ Pdcd1+), and (L/M) increase in effector CD8 T cells (CD45+ CD11B+ CD11C−LY6G− LY6C+). One-way ANOVA followed by Tukey’s Honest Significant Difference (HSD) test for pairwise comparisons).

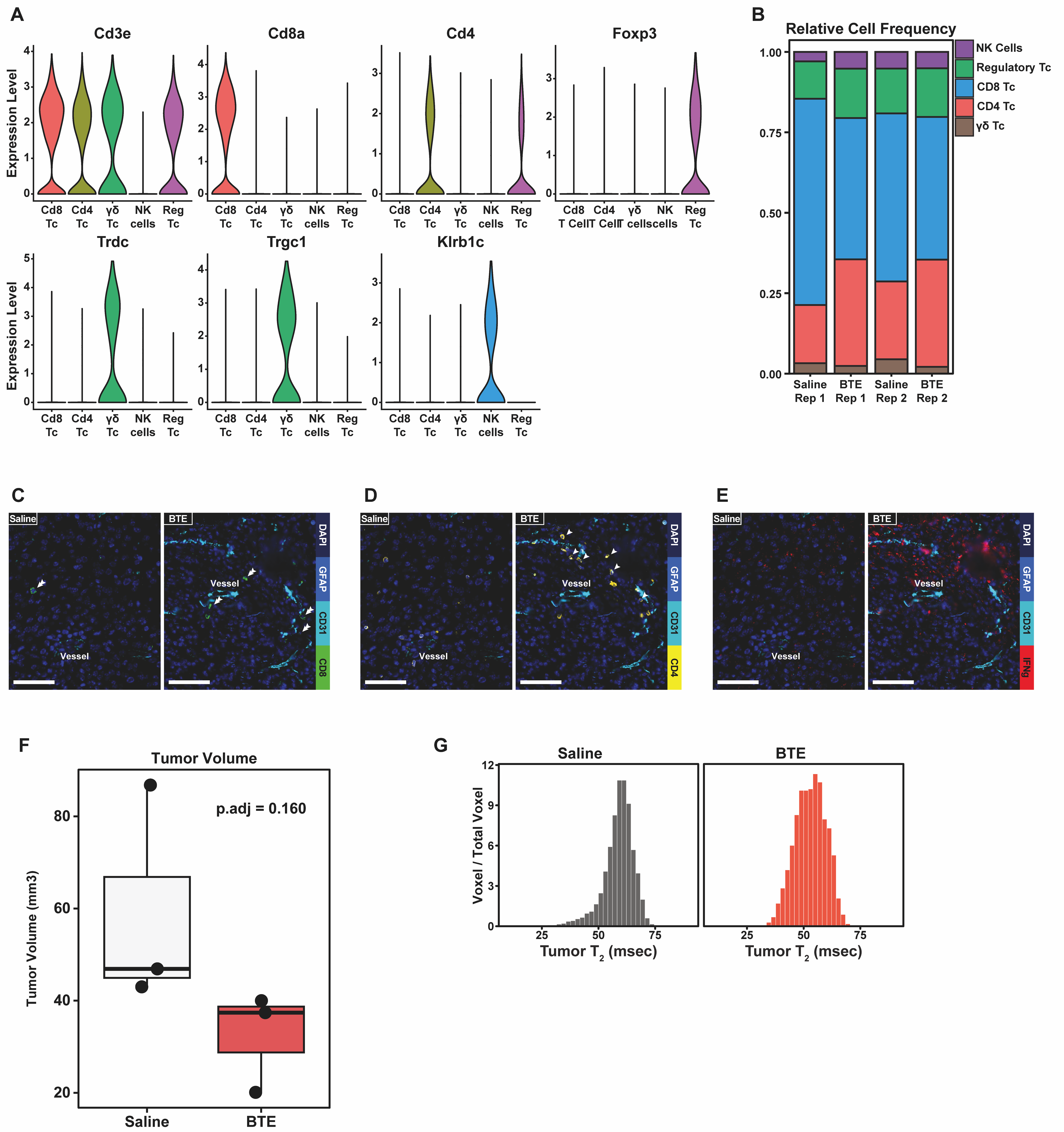

**Supplementary Figure 3. Identifying T and NK cell subcluster markers from single-cell RNA seq and tumor volume from MRI.** (*A*) Relative expression of Cd3e, Cd8a, Cd4, Foxp3, Trdc, Trgc1, and Klrb1c used as identifying markers of T and NK cell subcluster of single cell RNA seq. (*B*) Relative cell frequency of CD8 T cells (blue), CD4 T cells (red), regulatory T cells (green), γδ T cells (brown), and NK cells (purple) separated by sample treatment. (*C*) Box plot of tumor volume from MRI of BTE (n=3) and Saline (n=3) treated animals. (*D*) Histogram of voxel T2 MRI images separated by BTE and Saline treatment groups.
